# Co-circulation of multiple influenza A variants in swine harboring genes from seasonal human and swine influenza viruses

**DOI:** 10.1101/2020.07.28.225706

**Authors:** Pia Ryt-Hansen, Jesper Schak Krog, Solvej Østergaard Breum, Charlotte Kristiane Hjulsager, Anders Gorm Pedersen, Ramona Trebbien, Lars Erik Larsen

**Affiliations:** Technical University of Denmark, National Veterinary Institute, Kemitorvet building 204, 2700 Kgs. Lyngby, Denmark; University of Copenhagen, Department of Health Sciences, Institute for Animal and Veterinary Sciences 1870 Frederiksberg C, Denmark; Statens Serum Institut, Artillerivej 5, 2300 Copenhagen S, Denmark; Department of Health Technology, Section for Bioinformatics, Technical University of Denmark, Kemitorvet Building 204, DK-2800 Kongens Lyngby, Denmark

## Abstract

Since the influenza pandemic in 2009, there has been an increased focus on swine influenza A virus (swIAV) surveillance. This paper describes the results of the surveillance of swIAV in Danish swine from 2011 to 2018.

In total, 3800 submissions were received with a steady increase in swIAV positive submissions, reaching 56% in 2018. Ten different swIAV subtypes were detected. Full genome sequences were obtained from 129 swIAV positive samples. Altogether, 17 different circulating genotypes were identified including novel reassortants and subtypes harboring human seasonal IAV gene segments. The phylogenetic analysis revealed substantial genetic drift and also evidence of positive selection occurring mainly in antigenic sites of the hemagglutinin protein and confirmed the presence of a swine divergent cluster among the H1pdm09Nx viruses.

The results provide essential data for the control of swIAV in pigs and for early detection of novel swIAV strains with zoonotic potential.

## Introduction

Influenza A virus (swIAV) infection in swine causes respiratory disease, impairs the growth rate and increases the risk of secondary infections^1–3^. SwIAV is enzootic globally and multiple subtypes and lineages have been identified^4^. The influenza A virus genome consists of eight distinct gene segments and subtypes are assigned by characterizing the two surface glycoproteins hemagglutinin (HA) and neuraminidase (NA)^5^.

Pigs are infected by the same subtypes as humans, including H1N1, H1N2 and H3N2^6^. The transmission of H1N1 avian influenza A virus (IAV) to swine in the 1970s created the H1N1 Eurasian swine lineage also called “avian-like swine H1N1” (1.C lineage^7^) circulating in Europe and Asia^8^. An H3N2 influenza virus related to a human strain from 1973 started to circulate in the European pig populations in 1984. In the mid-1980s, a reassortment between the avian-like swine H1N1 and H3N2 human virus resulted in a human-like reassortant swine “H3N2sw” that became established in European swine^9,10^. In 1994, a H1N2 reassortant (1.B lineage) comprising an HA gene from human seasonal H1N1, an NA gene from H3N2sw and internal genes originating from avian-like swine H1N1 was first identified in the United Kingdom and subsequently detected in many European countries^11^. This swIAV lineage is also known as European human-like “H1huN2”. However, this subtype has never been detected in Danish pigs. In the beginning of the 2000s, a new “H1N2dk” reassortant virus was identified in Danish pigs^12^. This H1N2dk virus comprised an avian-like swine HA gene and an NA from contemporary, circulating H3N2sw and has since been identified in several European countries^13,14,15,16^.

In 2009, a novel IAV identified as pandemic H1N1/2009 strain of influenza A (1A.3.3.2 lineage - H1N1pdm09) spread rapidly among humans worldwide. The H1N1pdm09 virus is a reassortant, which obtained most of its gene segments from the triple-reassortant swIAV circulating in North American swine, its NA and matrix (M) gene segments from the Eurasian avian-like swine H1N1 lineage^17,18^ and had its origin in the Mexican swine population^19^. Soon after the virus began to spread globally in humans, its introduction into the swine population was noticed in several countries^15,20–22^. In transmission experiments, the high susceptibility of pigs to H1N1pdm09 infection was confirmed as well as an efficient pig-to-pig transmission^23^. This instantly raised concerns about the possible generation of new reassortants between H1N1pdm09 virus and circulating swIAV lineages, which soon after was indeed found to have occurred in several countries, including Denmark^15,24–27^. Consequently, there is a risk for the development of novel and more virulent progeny virus capable of infecting humans.

Surveillance of swIAV in pigs concerns both animal and public health. For animal health, the documentation of enzootic and new emerging swIAV and their ecology is important for control of disease and to ensure the use of adequate diagnostic tools. From a public health point-of-view, the results are important for risk assessments of emerging IAV, resistance to antiviral drugs or increased pathogenicity as well as pandemic preparedness. Here we report the results of a passive surveillance program of swIAV conducted in Denmark from 2011 to 2018, including data on intensive subtyping and genetic characterization of swIAV positive submissions.

## Results

### Field samples

The total number of submissions received for swIAV diagnostics from pigs with acute respiratory disease in the years 2011 to 2018 fluctuated over the years, with a peak in 2015 (Fig. 1). In total, 3800 submissions were received over eight years. The pattern of monthly submissions was very similar from year to year showing a peak in the number of submissions from October to March (autumn and winter months) (Fig. 1). When comparing the number of swine herds submitting samples for swIAV diagnostics each year (n=276-488) with the total number of swine herds present in Denmark the same year (n=2741-4529)^28^, it was evident that 6-15% of the Danish swine herds were included in the surveillance with a steady increase over the years.

**Fig. 1.**
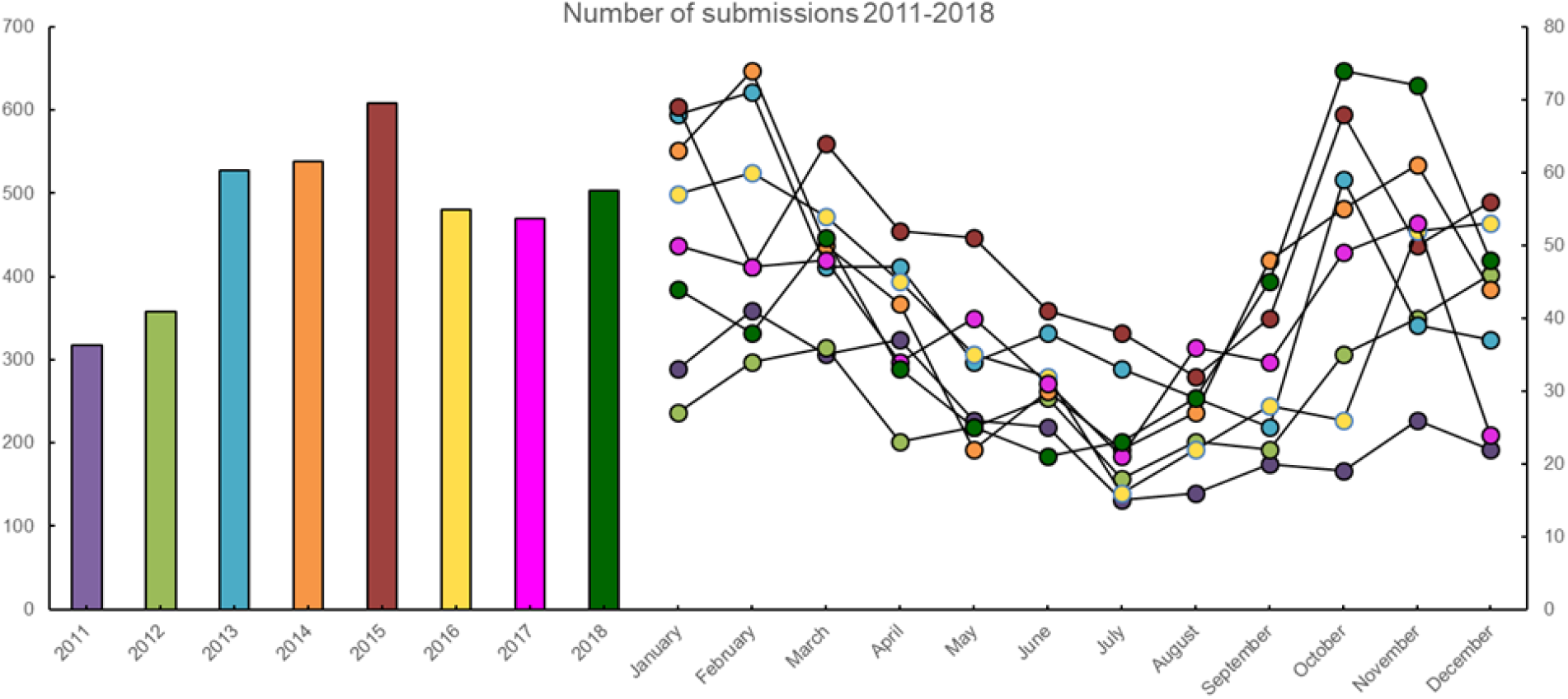
The annual and monthly number of submissions received from Danish pigs with acute respiratory disease in the years 2011 to 2018.

### SwIAV positive samples

In 2011, the first year of the surveillance, 36 % of the total submissions contained at least one positive sample. In the following five years (2012-2016), the percentage of swIAV positive submissions was stable ranging between 44-47 %, but, an increase in the percentage of swIAV positive submission was observed over the last two years, reaching 56 % in 2018 (Fig. 2). However, it should be noted that the average number of samples per submission was higher in 2018, with an average of 2.9 samples per submission compared to 2-2.3 the previous years (2011-2017) (data not shown). The monthly distribution of swIAV positive submissions was fluctuating, but no consistent seasonal variations were observed (Supplementary figure 1). The average monthly percentage of positive submission over the eight years ranged from 42.6-51.8 % with the highest average percentages in April, September and December. There was no significant difference between the average percentage of positive submissions between the different months, with the exception of April (average percentage of positive submissions = 51.6 %, SD: 6.5) and August (average percentage of positive submissions = 42.6 %, SD: 8.9) (p=0.04).

**Fig. 2.**
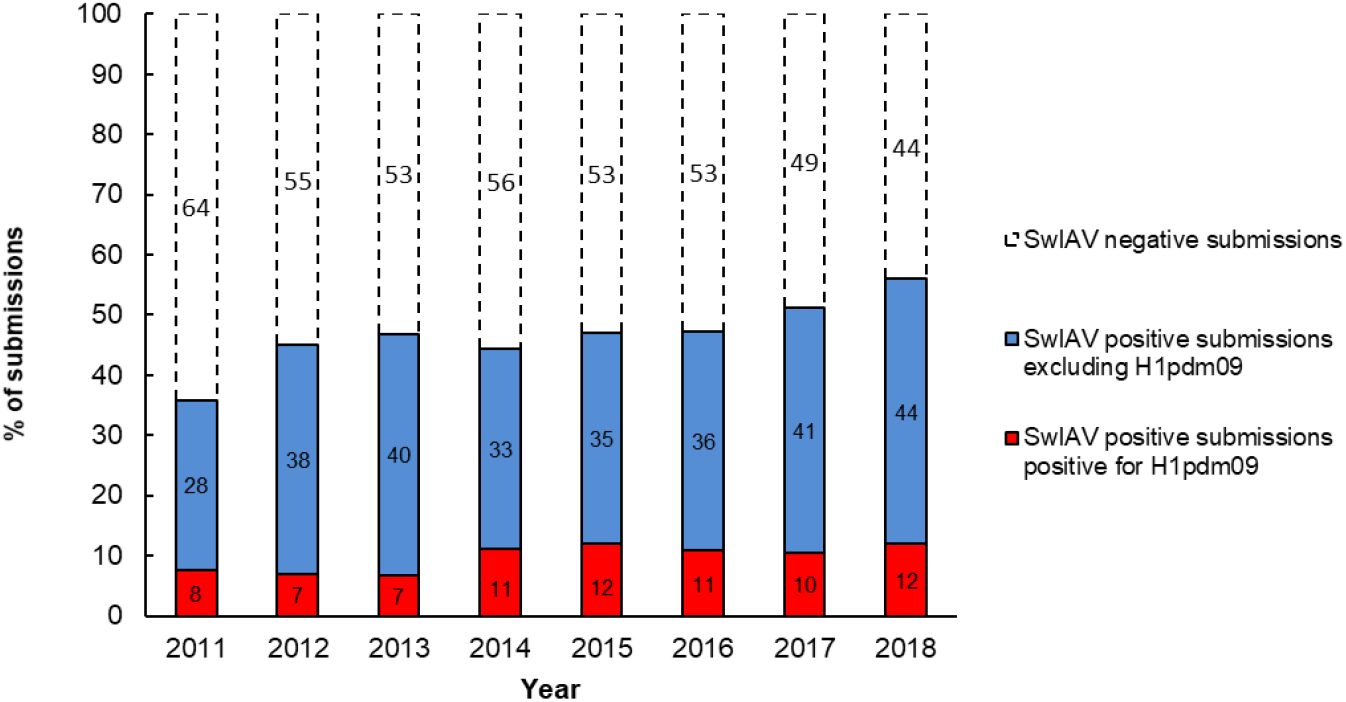
The percentages of submissions testing positive and negative for influenza A virus and the proportion of positive submission testing positive for H1pdm09 by real time RT-PCR from 2011-2018.

### Test for the HA gene of H1N1pdm09 origin by specific real time PCR

Due to the global spread of H1N1pmd09 virus in humans, it was decided in 2011 to test all swIAV positive samples from Danish pigs specifically for the presence of the HA gene of H1N1pdm09 origin (H1pdm09). In 2011, 21 % of the swIAV positive submissions, tested positive for H1pdm09. However, in the two following years the percentage decreased to 14-16 % of the swIAV positive submissions. This decrease reverted in 2014, where a marked increase was observed, and since then, the proportion of H1pdm09 remained at a stable level, ranging between 20-26 % of the swIAV positive submissions (Fig. 3). On average over the eight years, H1pdm09 positive submissions constituted 15-25% of the monthly swIAV positive submissions (Supplementary figure 1). The months with the highest proportion of H1pdm09 positive submissions were February, March and July. However, no significant differences (p value >0.05) in the average proportions of H1pdm09 positive submissions between the different months were observed.

**Fig. 3.**
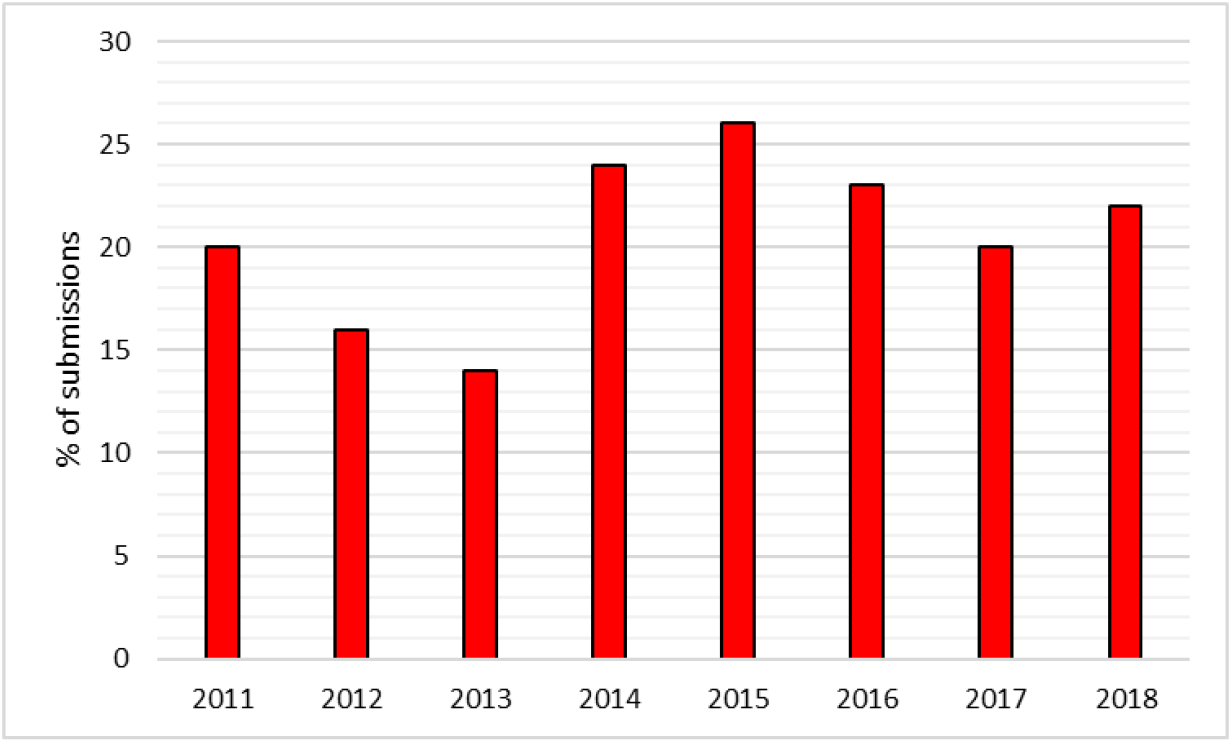
The percentages of the swIAV positive submissions testing positive in the screening for H1pdm09 by real time RT-PCR from 2011-2018.

### SwIAV subtypes

During the years 2011-2014, 33-48 % of the swIAV positive submissions were subtyped by partial sequencing of the HA and NA surface genes. From 2015 and onwards, the swIAV positive submissions was subtyped by multiplex RT-PCR and/or by Fluidigm, resulting in an increase of successfully subtyped submissions to 61-77 % of the total number of swIAV positive submissions (data not shown).

The most prevalent subtype identified in the swIAV positive submissions during the eight years (2011-2018), was H1N2dk. The proportion of H1N2dk increased steadily from 42 % of the subtyped submissions in 2012 to 69 % of the subtyped submissions in 2018. In contrast, the avian-like swine H1N1, which was highly prevalent in 2011-2012 representing between 30-37 % of the subtyped submissions, decreased markedly since then, only representing 5% of the swIAV subtyped submissions in 2018 (Fig. 4).

**Fig. 4.**
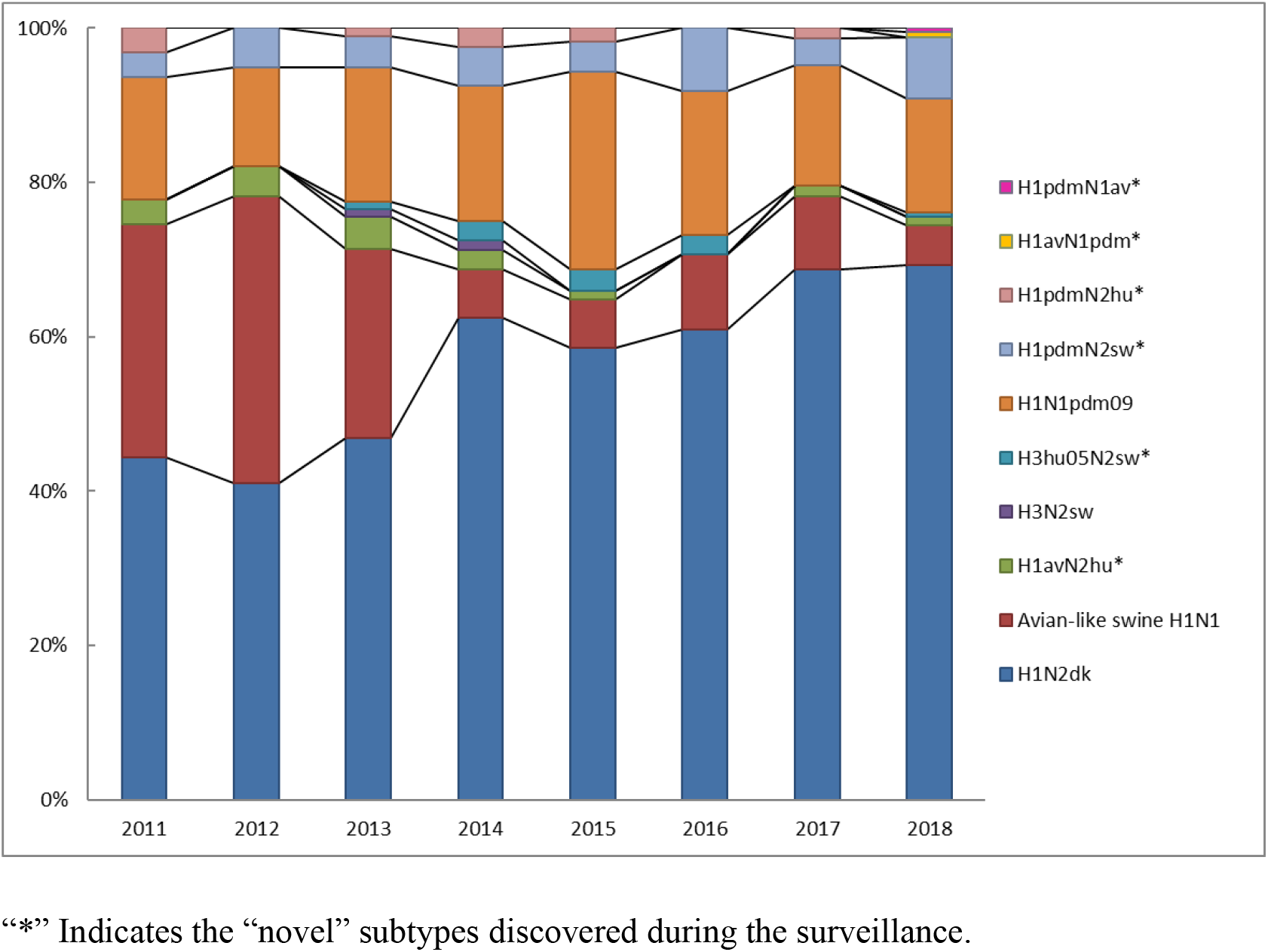
Subtype distribution shown as the percentage of total number of subtyped submissions from 2011-2018. “*” Indicates the “novel” subtypes discovered during the surveillance.

The proportion of H1N1pdm09 was relatively stable from 2011-2018 representing approx. 16 % of the subtyped submissions. However, in 2015 a marked increase was observed when the H1N1pdm09 was detected in 25.6 % of the subtyped submissions. Several reassortants containing either the HA or the NA gene of the H1N1pdm09 subtype, were identified. The most prevalent of these reassortants was the “H1pdmN2sw”, which combined the HA gene of the H1N1pdm09 subtype and the NA gene of the H1N2dk subtype. This subtype was detected for the first time in Denmark in 2011, and since then, the proportion of this subtype remained relatively stable constituting around 4 % of the subtyped submissions each year. However, in the years 2016 and 2018 a doubling in prevalence of H1pdmN2sw was observed. Another reassortant, also detected for the first time in 2011, was the “H1pdmN2hu”, which contained an HA gene of H1N1pdm09 origin, and an NA gene derived from the human seasonal flu circulating in the 90’s. This subtype was identified with low prevalence (1-3.2 %) from 2011-2017, but was interestingly not detected, in the years where the prevalence of H1pdmN2sw peaked (2016 and 2018). In 2018, two novel swIAV subtypes were identified, both including one surface gene of H1N1pdm09 origin. One was termed “H1avN1pdm” and had an avian-like swine HA gene and an NA gene of H1N1pdm09 origin. The other novel subtype was termed “H1pdmN1av” and carried an HA gene of H1N1pdm09 origin and an NA gene derived from the avian-like swine H1N1 (Fig. 4).

The swine-adapted reassortant H3N2sw was detected in a few samples in 2013-2014, but was not detected in 2015-2018. However, another H3N2 reassortant “H3hu05N2sw”, containing an HA gene of human seasonal origin from 2005 and an NA gene of the H1N2dk subtype, has been detected each year since 2013, with the exception of 2017^26^. Another reassortant, containing the N2 gene of the human seasonal H3N2 subtype, was detected for the first time in Denmark in 2011 and was termed “H1avN2hu”^29^. This subtype carried an avian-like swine HA gene and an NA gene derived from the human seasonal flu circulating in the 90’s. The H1avN2hu subtype has been detected each year, with the exception of 2016 (Fig. 4).

In summary, six novel swIAV reassortant subtypes (H1pdmN2sw, H1pdmN2hu, H1pdmN1av, H1avN1pdm, H3hu05N2sw and H1avN2hu) were discovered through the Danish surveillance of swIAV from 2011-2018 (Fig. 4). However, the diversity of circulating strains is even more complex, when all gene segments are included in the analyses as described below.

### Full genome sequencing

In total, 128 full genome sequences of swIAV isolated between 2013-2018 were uploaded in GenBank with the following accession numbers: MT666225 - MT667233. The accessions numbers, corresponding sample IDs and information on the lineage of each gene segments are summarized in Supplementary table 2. The characteristics of the H3hu05N2sw subtype (isolate 2014_15164_1p1_H3hu05N2sw accession number: EPI_ISL_247092) has previously been described^26^, but was also included in the analysis of the H3hu genes of this study.

### Hemagglutinin gene characterization

In total, 78 H1av, 48 H1pdm09 and three H3hu05 full-length HA sequences were obtained and analyzed separately according to the lineage.

The H1av nucleotide sequences were fairly diverse, with an average pairwise sequence difference per site (pi) of 0.099, SE: 0.0005. Phylogenetic trees constructed either with or without clock-models (Supplementary figure 2, Figure 5), did not display the imbalanced, ladder-like structure typical for influenza trees. The tree contained several clusters and one cluster (Cluster 6, Figure 5) that was dominated by H1avNx strains carrying a complete internal gene cassette of H1N1pdm09 origin. There was low correlation between sampling time and genetic divergence (Table 2).

**Fig. 5.**
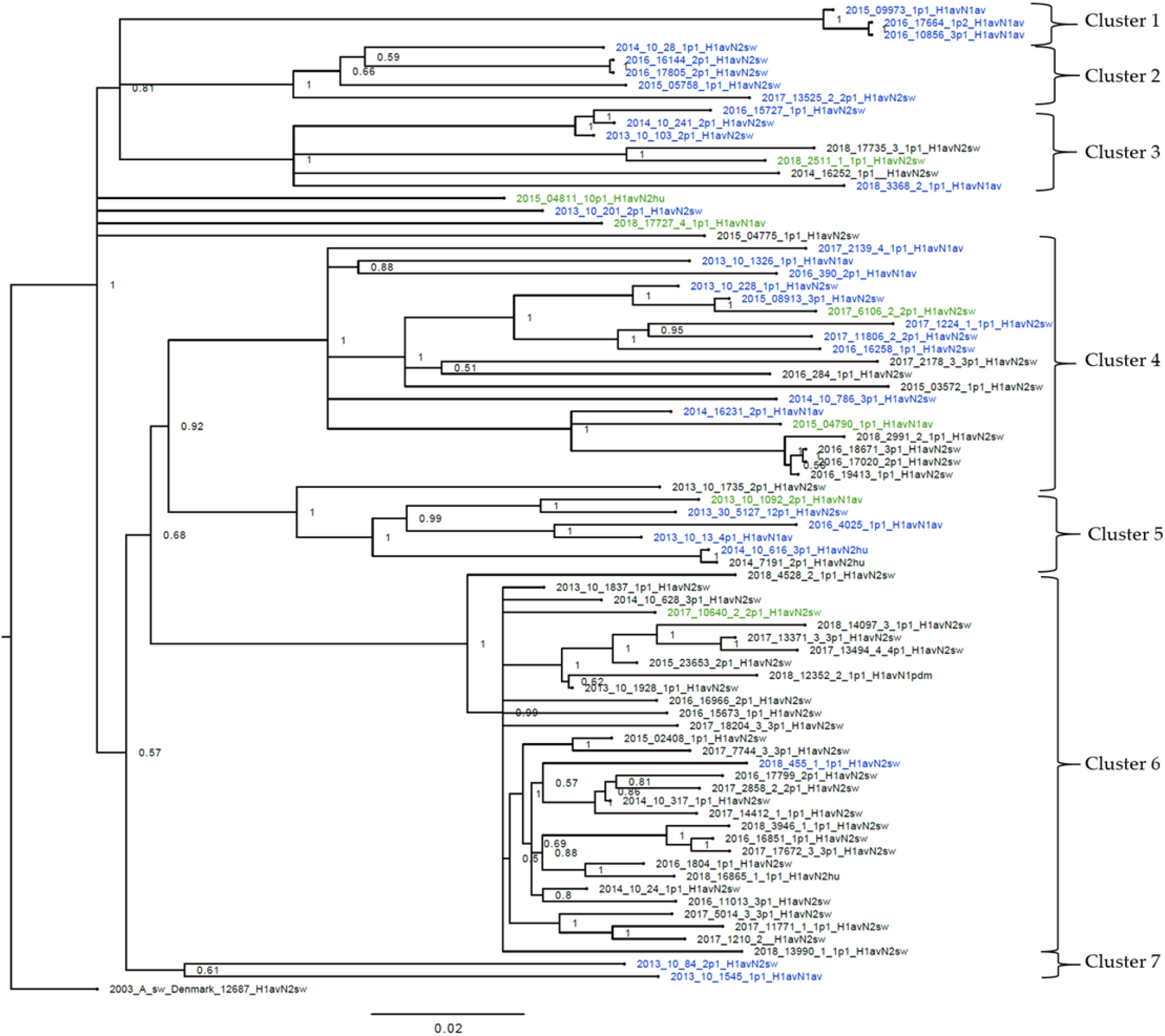
Bayesian phylogenetic tree of the H1av nucleotide sequences. Node labels represent posterior probabilities. “2003_A_sw_Denmark_12687” is the outgroup. The taxon includes the year wherefrom the sample were obtained, the sample ID and subtype. A blue taxon indicates that the internal gene cassette is of avian-like swine origin, a green internal gene cassette indicates that the internal cassette has a mix of avian-like swine and H1N1pdm genes and a black taxon indicates that the internal gene cassette is of H1N1pdm09 origin.

**Table 1.**
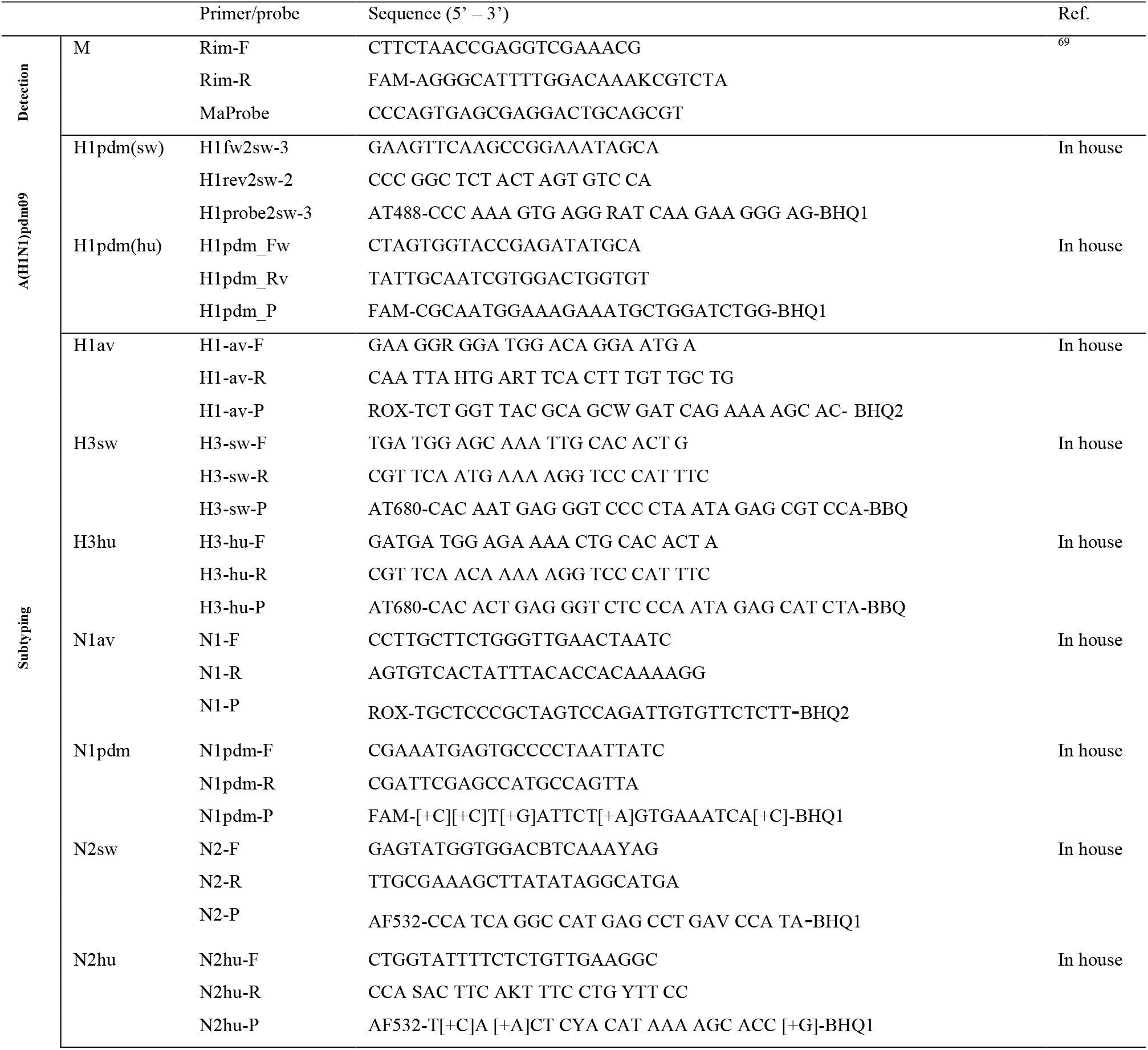
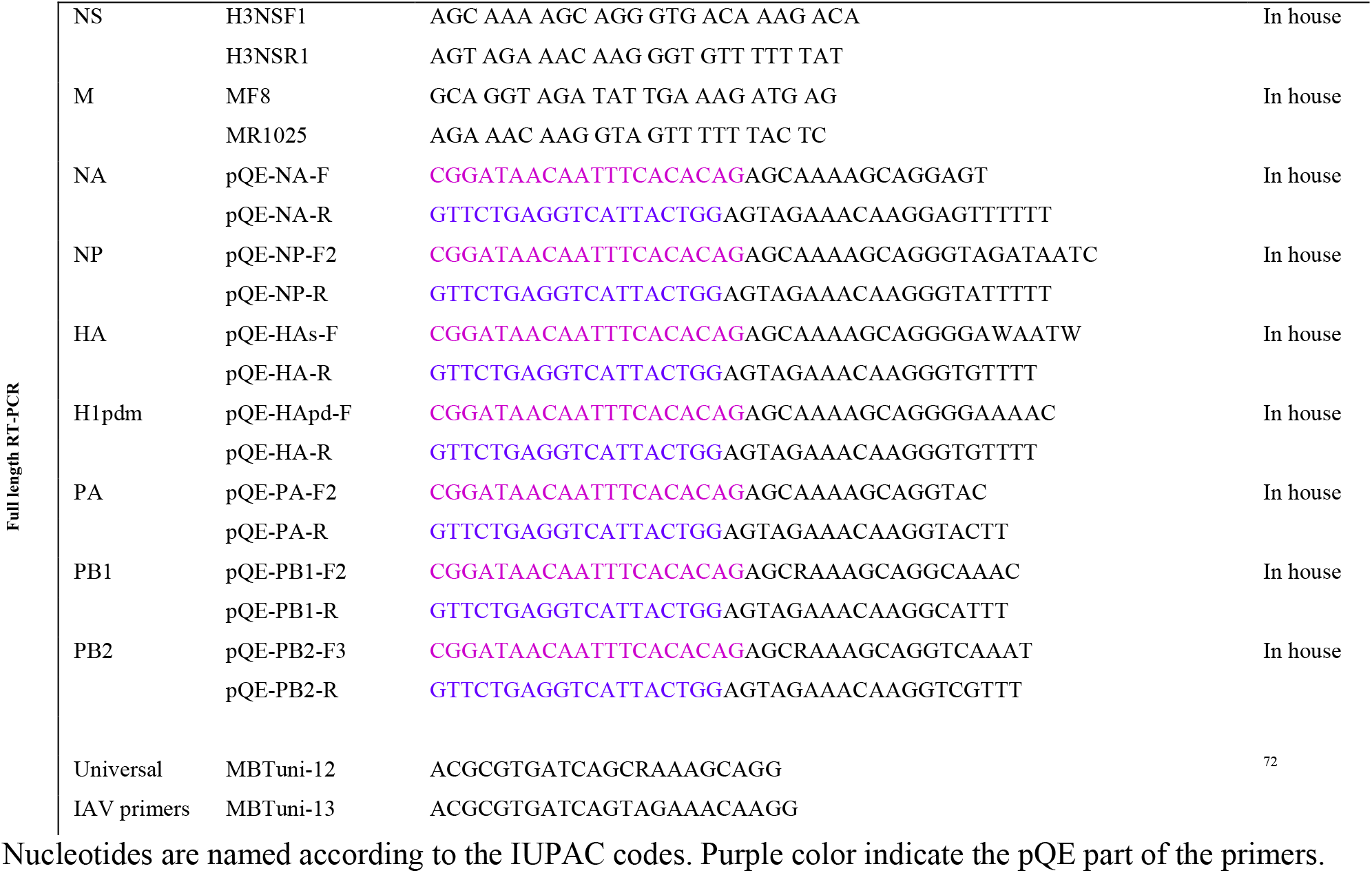
Primers and probes used for detection, subtyping and full genome sequencing of swIAV

**Table 2.**
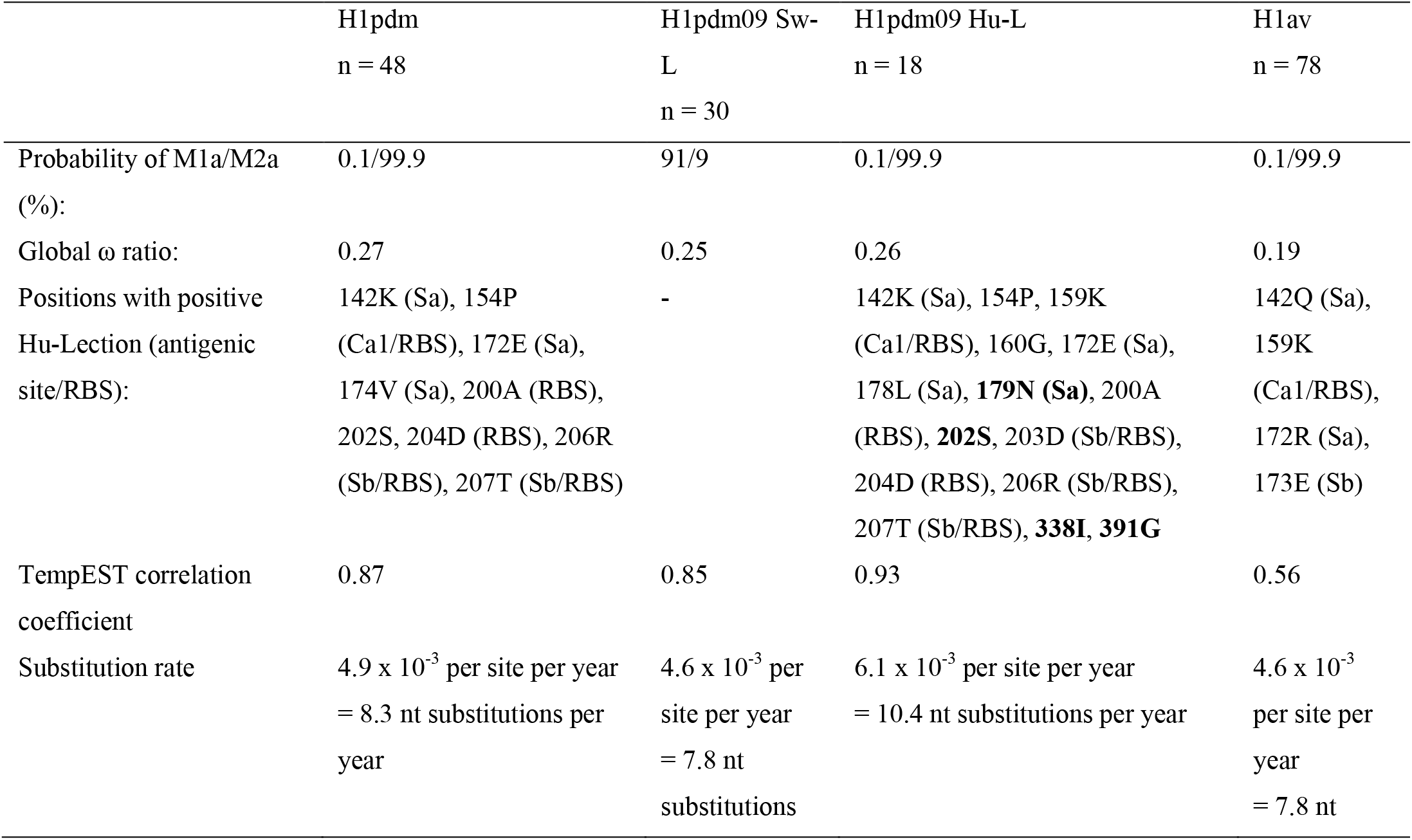
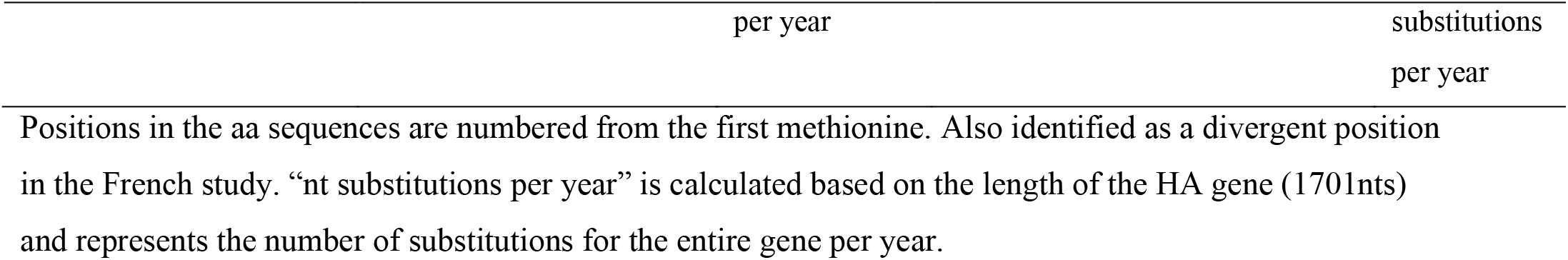
Results of the evolutionary analysis of the HA gene/protein of the H1av and the H1pdm09 lineages:

|  | H1pdm<br>n = 48 | H1pdm09 Sw-L<br>n = 30 | H1pdm09 Hu-L<br>n = 18 | H1av<br>n = 78 |
| --- | --- | --- | --- | --- |
| Probability of M1a/M2a<br>(%): | 0.1/99.9 | 91/9 | 0.1/99.9 | 0.1/99.9 |
| Global $\omega$ ratio: | 0.27 | 0.25 | 0.26 | 0.19 |
| Positions with positive<br>Hu-Lection (antigenic<br>site/RBS): | 142K (Sa), 154P<br>(Ca1/RBS), 172E (Sa),<br>174V (Sa), 200A (RBS),<br>202S, 204D (RBS), 206R<br>(Sb/RBS), 207T (Sb/RBS) | - | 142K (Sa), 154P, 159K<br>(Ca1/RBS), 160G, 172E (Sa),<br>178L (Sa), **179N (Sa)**, 200A<br>(RBS), **202S**, 203D (Sb/RBS),<br>204D (RBS), 206R (Sb/RBS),<br>207T (Sb/RBS), **338I**, **391G** | 142Q (Sa),<br>159K<br>(Ca1/RBS),<br>172R (Sa),<br>173E (Sb) |
| TempEST correlation<br>coefficient | 0.87 | 0.85 | 0.93 | 0.56 |
| Substitution rate | 4.9 x 10<sup>-3</sup> per site per year<br>= 8.3 nt substitutions per<br>year | 4.6 x 10<sup>-3</sup> per<br>site per year<br>= 7.8 nt<br>substitutions | 6.1 x 10<sup>-3</sup> per site per year<br>= 10.4 nt substitutions per year | 4.6 x 10<sup>-3</sup><br>per site per<br>year<br>= 7.8 nt |

Analysis using CODEML indicated strong evidence for positive selection among the H1av sequences. Specifically, the dN/dS ratios for individual codons under the M2a model strongly suggested the presence of positive selection in four positions, all located in the globular head of the HA protein and all in previously defined antigenic sites (Table 2).

The H1pdm09 nucleotide sequences had a lower nucleotide diversity: pi = 0.043, SE 0.0005. Both the clock and non-clock trees for H1pdm09 sequences isolated from Danish pigs, showed that 30 of the sequences were located in a well-defined cluster (Cluster 1; Fig. 6 and Supplementary figure 3), with the remaining 18 sequences branching out basally to this cluster (Fig. 6 and Supplementary figure 3). The 30 H1pdm09 sequences of Cluster 1 were collected between 2015-2018, whereas the 18 H1pdm09 sequences outside this cluster were collected between 2013-2017 (Fig. 6). The strict molecular clock tree and the TempEst analysis of all the Danish H1pdm09 sequences suggested that the sequences evolved according to time with stable substitution rate of 4.9 × 10^−3^ per site per year (Fig. 6 and Table 2). The diversion into the Cluster 1 appeared to have occurred around 2011, however the most recent common ancestor for the sequences in Cluster 1 was dated around 2014 (Fig. 6). In the phylogenetic tree that also included representative swine and human seasonal H1pdm09 sequences, it was found that Cluster 1 only contained swine derived H1pdm09 sequences, while sequences outside of this cluster was a mix of swine and human seasonal H1pdm09 sequences. Cluster 1 was therefore termed the swine like “Sw-L cluster” (Fig. 7 – Sw-L cluster – taxon suffix Sw-L). The remaining 18 Danish swIAV sequences were termed human like “Hu-L” H1pdm09 sequences (Fig. 7 - taxon suffix Hu-L).

**Fig. 6.**
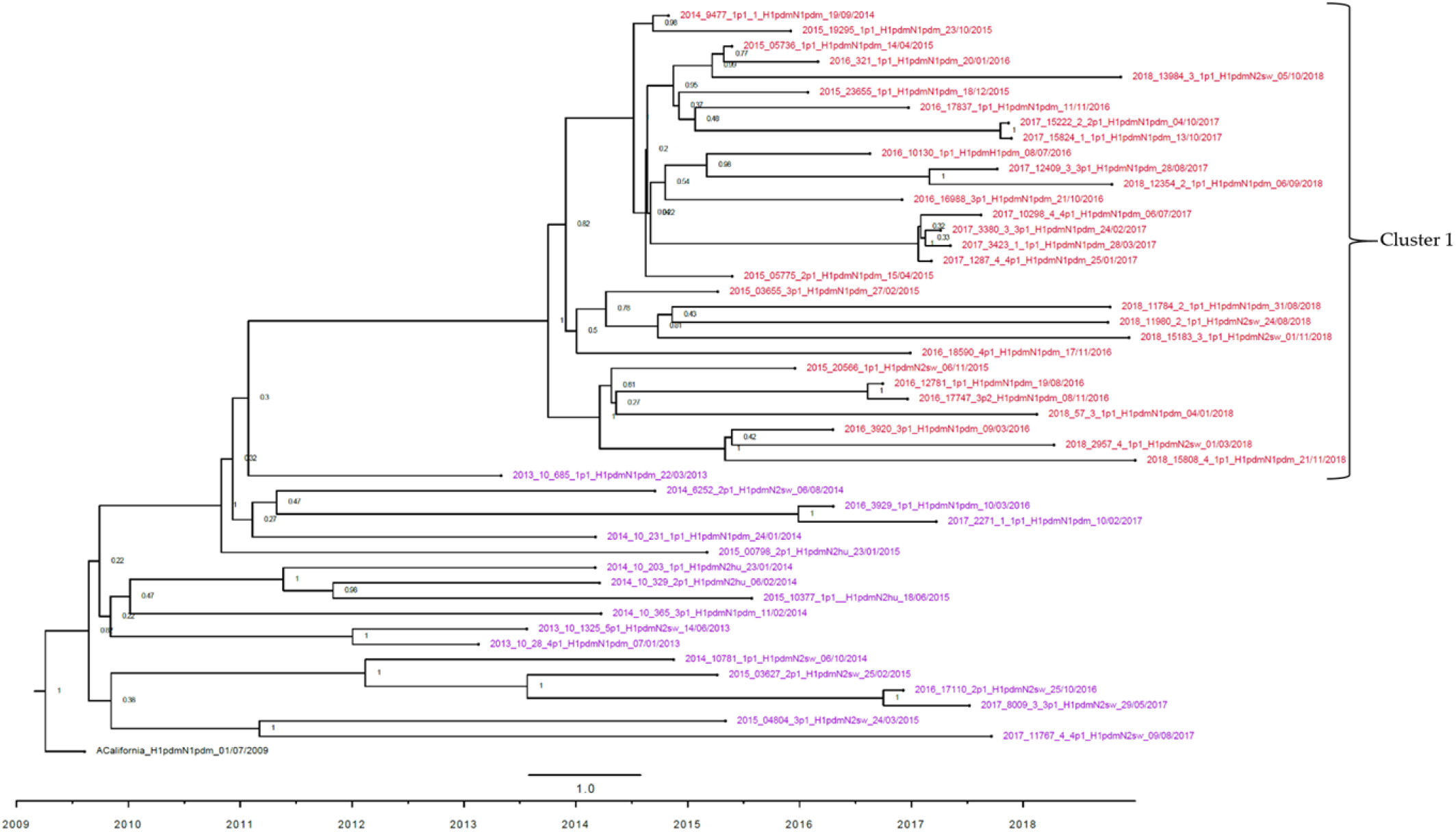
Strict molecular clock tree of the Danish H1pdm09 sequences. Node labels represent posterior probabilities. The x-axis represents the time in years, and each tick indicates half a year. The taxon includes the year wherefrom the sample were obtained, the sample ID and subtype. A red taxon indicates samples included in “Cluster 1” and a purple taxon indicates samples located outside “Cluster 1”.

**Fig. 7.**
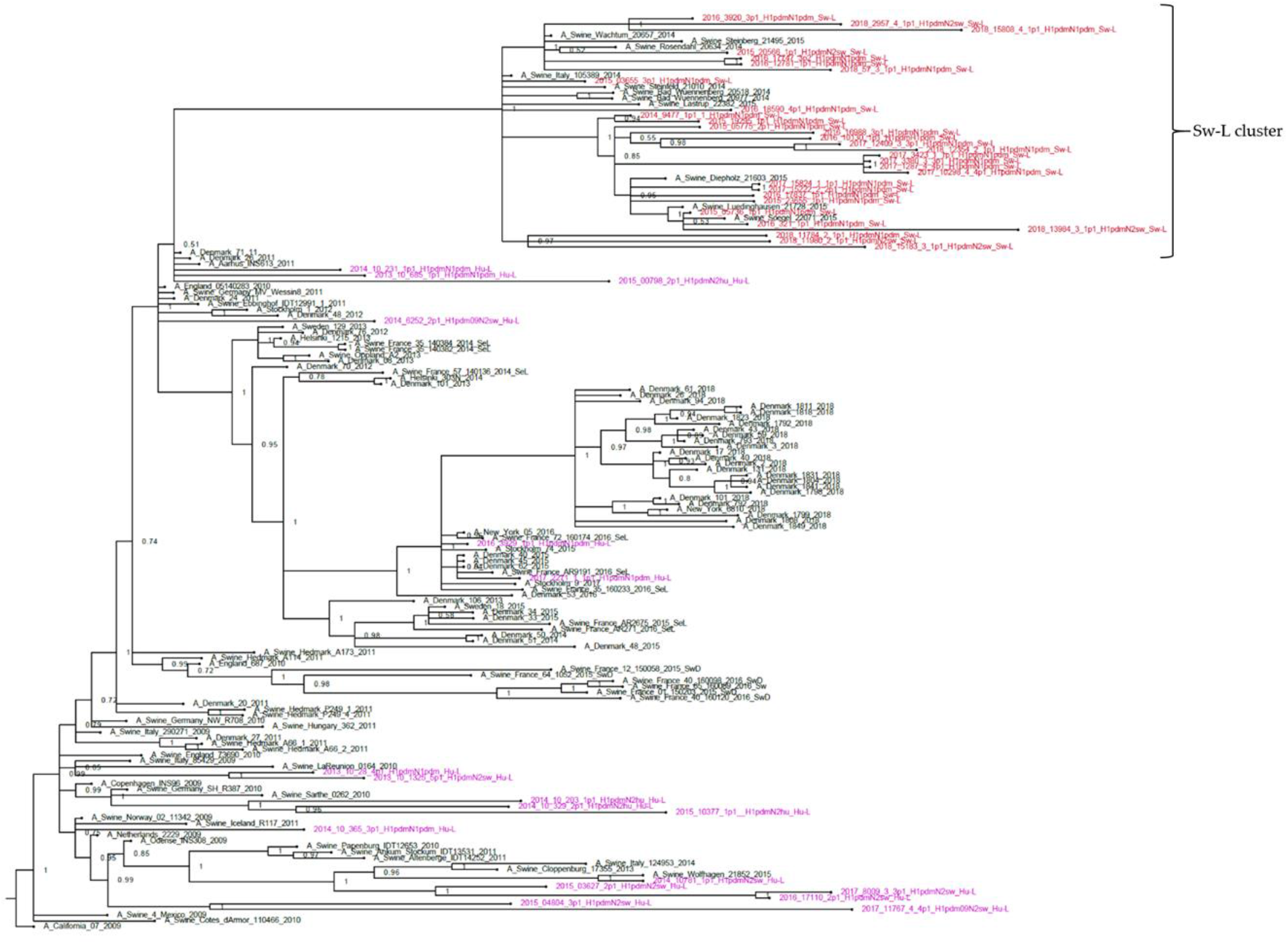
Bayesian phylogenetic tree with of H1pdm09 sequences including reference strains Node labels represent posterior probabilities. A_California_2009 is the outgroup. The reference sequences are named according to their given name in NCBI GenBank or GISAID and year of isolation. In addition, the French swine derived sequences have the suffix “SwD” or “SeL”. The red taxons with the suffix “Sw-L” correspond to the Danish sequences of Cluster 1 - Fig. 6 and is now included in the “Sw-L” cluster. The purple taxons correspond to the purple taxons of Fig. 6 and have been given the suffix “Hu-L”.

The initial analysis of the Danish H1pdm09 aa sequences revealed a total of 20 aa positions that differed between the Danish Sw-L and the Danish Hu-L sequences (Table 3) and seven of these 20 aa differences were specific, meaning that all the 30 Danish Sw-L aa sequences had a different aa compared to all of the 18 Danish Hu-L aa sequences (Bold positions in Table 3). Thirteen of the 20 aa residues defining the Sw-L protein sequences were located either in previously defined antigenic sites (Ca and Sb) or the receptor binding site (RBS). Six of these were among the seven “unique” Sw-L positions (Table 3). Subsequently, the 20 aa residues were compared among all the sequences included in the phylogenetic tree of Fig 7. These reference aa sequences were divided into three groups; one containing the foreign (non-Danish) swine H1pdm09 sequences (n=11) included in the “Sw-L cluster”, one containing the European swine H1pdm09 sequences located outside the Sw-L cluster (n=42) and one containing human seasonal H1pdm09 sequences (n=59) (Table 3). Interestingly, all the 11 foreign swine H1pdm09 aa sequences included in the Sw-L cluster, shared exactly the same aa residues as the Danish Sw-L sequences. Similarly, the majority of the European swine- and human seasonal H1pdm09 aa sequences located outside the Sw-L cluster carried residues similar to the Danish Hu-L aa sequences, and were different from the sequences included in the Sw-L cluster (Table 3). Finally, no unique swine or human residues were revealed when all H1pdm09 proteins derived from swine were compared to the H1pdm09 proteins derived from human seasonal H1pdm09 viruses. Nonetheless, at position 273, significantly more swine H1pdm09 proteins (91 %) carried an A compared to the human seasonal H1pdm09 proteins (27 %) (p = < 0.05). In summary, the H1pdm09 proteins derived from Danish pigs were divided into two groups containing “Sw-L” and “Hu-L” sequences, which were separated by 20 aa differences mainly located in antigenic sites or the RBS. The Sw-L cluster still clustered separately when swine- and human-derived H1pdm09 reference sequences were included in the alignment, but 11 additional German and Italian H1pdm09 swine-derived sequences were also part of this cluster.

**Table 3.**
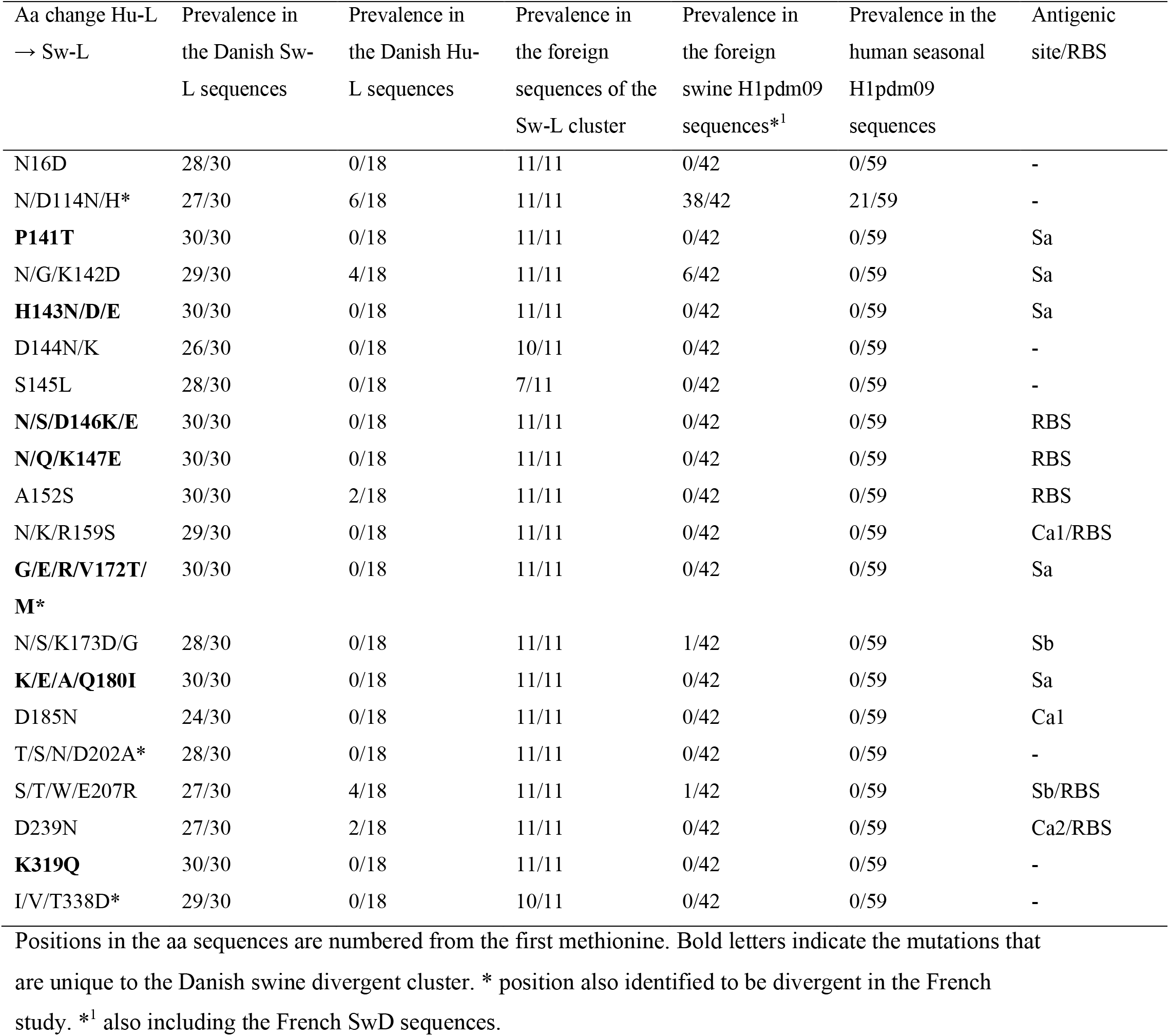
Mutations of the H1pdm09 defining the Danish swine divergent clusters in relation to the seasonal-like H1pdm09 sequences.

The CODEML analysis for determining the best fitting substitution model revealed that the M2a model fitted the Danish H1pdm09 sequences significantly better than the M1a model, providing strong evidence for positive selection occurring in the HA protein. Moreover, the dN/dS ratios for individual codons under the M2a model strongly suggested the presence of positive selection in nine aa positions all situated in the globular part of the HA protein and eight located specifically in antigenic sites or in the RBS. The same analysis was repeated on the Danish Sw-L and Hu-L sequences, separately. Interestingly, these analyses revealed that positive selection did indeed occur in the Danish Hu-L sequences, as the M2a model fitted the sequences significantly better than the M1a model. Additionally, the dN/dS ratios for individual codons under the M2a model strongly suggested the presence of positive selection in 15 different aa positions, including ten positions located in antigenic sites or the RBS. Conversely, model M1a fitted the Sw-L sequences significantly better, suggesting that no positive selection occurred among these sequences. The strict molecular clock and TempEst analysis were also repeated for the Danish Sw-L and Hu-L sequences separately. Interestingly, the Hu-L sequences had a higher substitution rate and also showed a higher correlation coefficient in the TempEst analysis compared to the Sw-L sequences (Table 3). In summary, when analyzing all the H1pdm09 sequences as a whole, positive selection was evident among the sequences. However, when dividing the H1pdm09 sequences into the Sw-L and Hu-L groups, positive selection was only evident among the Hu-L H1pdm09 sequences and these sequences also showed a higher substitution rate compared to the Sw-L sequences.

Additionally, differences in N-linked and O-linked glycosylation sites between the Sw-L and Hu-L H1pdm09 proteins were examined. The results revealed that all proteins of both the Sw-L and Hu-L samples were predicted to be N-glycosylated at position 28, 40, 304 and 557 (numbering from the first methionine). In addition, 3/18 Hu-L H1pdm09 proteins were predicted to be N-glycosylated at position 136, which is in the vicinity of the RBS. Significantly more Sw-L H1pdm09 proteins (80 %) had an O-linked glycosylation site at position 150 compared to the Hu-L H1pdm09 proteins (11%) (p=<0.05). Interestingly, position 150 is located in the RBS. Conversely, significantly more Hu-L H1pdm09 proteins (44 %) had a O-linked glycosylation site at position 145 compared to the Sw-L H1pdm09 proteins (3 %) (p=<0.05). Position 145 is also located in the close vicinity of the RBS.

Previously defined residues of the HA proteins regarded as important for host-adaptation, pathogenicity, receptor binding and virulence were examined and compared between subtypes carrying an HA protein of avian and H1N1pdm09 origin, respectively. The results can be visualized in Supplementary table 3.

The three H3 sequences obtained in this study, showed a low nucleotide diversity (pi) of 0.027, SE: 0.002. The closest human IAV match in NCBI GenBank for all of the three sequences was “A/Denmark/129/2005(H3N2)” with accession number EU103786. As only three sequences were obtained, no further phylogenetic or evolutionary analysis were performed.

### Neuraminidase characteristics

In total, 32 N1pdm, 14 N1av, 75 N2sw and 8 N2hu full-length NA sequences were obtained and analyzed separately according to the lineage.

The N1pdm nucleotide diversity (pi) was 0.029 SE: 0.0005. The majority of the sequences obtained between 2015-2018 were located in one cluster, whereas the oldest sequences (2013-2014) were located outside the cluster (Supplementary figure 4). The TempEst analysis showed a high correlation coefficient similar to that of the H1pdm09 sequences, indicating that the genetic divergence evolved according to time. The Beast analysis revealed a substitution rate of 3.9 × 10^−3^ per site per year. However, no evidence of positive selection was revealed (Table 4).

**Table 4.** Results of the evolutionary analysis of the NA gene of the N1pdm, N1av, N2sw and N2hu lineages:

|  | N1pdm | N1av | N2sw | N2hu |
| --- | --- | --- | --- | --- |
| Probability of M1a/M2a (%): | 90/10 | 73/27 | 88/12 | 88/12 |
| Global $\omega$ ratio: | 0.24 | 0.15 | 0.17 | 0.18 |
| Positions with positive Hu-Lecion: | - | - | - | - |
| TempEST correlation coefficient | 0.88 | 0.68 | 0.75 | 0.62 |
| Substitution rate (per site per year) | $3.9 \times 10^{-3}$ | $5.9 \times 10^{-3}$ | $4.4 \times 10^{-3}$ | N.A |

The N1av nucleotide diversity was 0.097, SE: 0.003 similar to the nucleotide diversity of the H1av nucleotide sequences. No clear clustering was observed in the Bayesian tree (Supplementary figure 5). The TempEST analysis showed a relative low correlation between the genetic divergence and time, and the Beast analysis revealed a substitution rate of 5.9 × 10^−3^ per site per year. No evidence of positive selection was observed (Table 4).

The N2sw nucleotide diversity was 0.08, SE: 0.0005 and the Bayesian analysis revealed six main clusters. Each cluster contained sequences dispersed over the majority of the surveillance period, suggesting no temporal clustering. Interestingly, one major cluster only contained HxN2sw from strains having a full or partial H1N1pdm09 internal gene cassette. Moreover, this cluster contained 28/30 of the same samples as Cluster 3 of the H1av sequences, which also clustered according to the origin of the internal gene cassette (Fig. 5 and Supplementary figure 6). The TempEst and Beast analysis revealed a low correlation between genetic divergence and time, and a substitution rate of 4.4 × 10^−3^ per site per year. As for the other NA lineages, no evidence of positive selection was observed (Table 4).

The eighth N2hu sequences showed a sequence diversity of 0.085, SE: 0.006 and despite the limited number of sequences, the Bayesian phylogenetic analysis revealed two main clusters; one containing sequences derived from subtypes containing a full avian internal gene cassette and one only containing sequences with a full or partial H1N1pdm09 internal cassette (Supplementary Figure 7). The TempEst analysis revealed a low correlation between genetic divergence and time, and the low number of sequences resulted in an overestimated substitution rate, which therefore was not included in the results. No evidence of positive selection was observed (Table 4).

All of NA sequences across the different lineages (n=129) were examined for specific aa changes encoding either neuraminidase resistance or increased virulence. However, none of the NA sequences had any of these aa changes.

### The internal gene cassette

In total, 17 different genotypes were identified in this study (Fig. 8). The subtypes H1N2dk, avian-like swine H1N1 and the H1avN2hu showed the highest number of diverse genotypes, whereas most subtypes including at least one surface gene of H1N1pdm09 origin had a complete internal gene cassette of H1N1pdm09 origin. Detailed information on the origin of all gene segments of all individual samples are listed in Supplementary table 2.

**Fig. 8.**
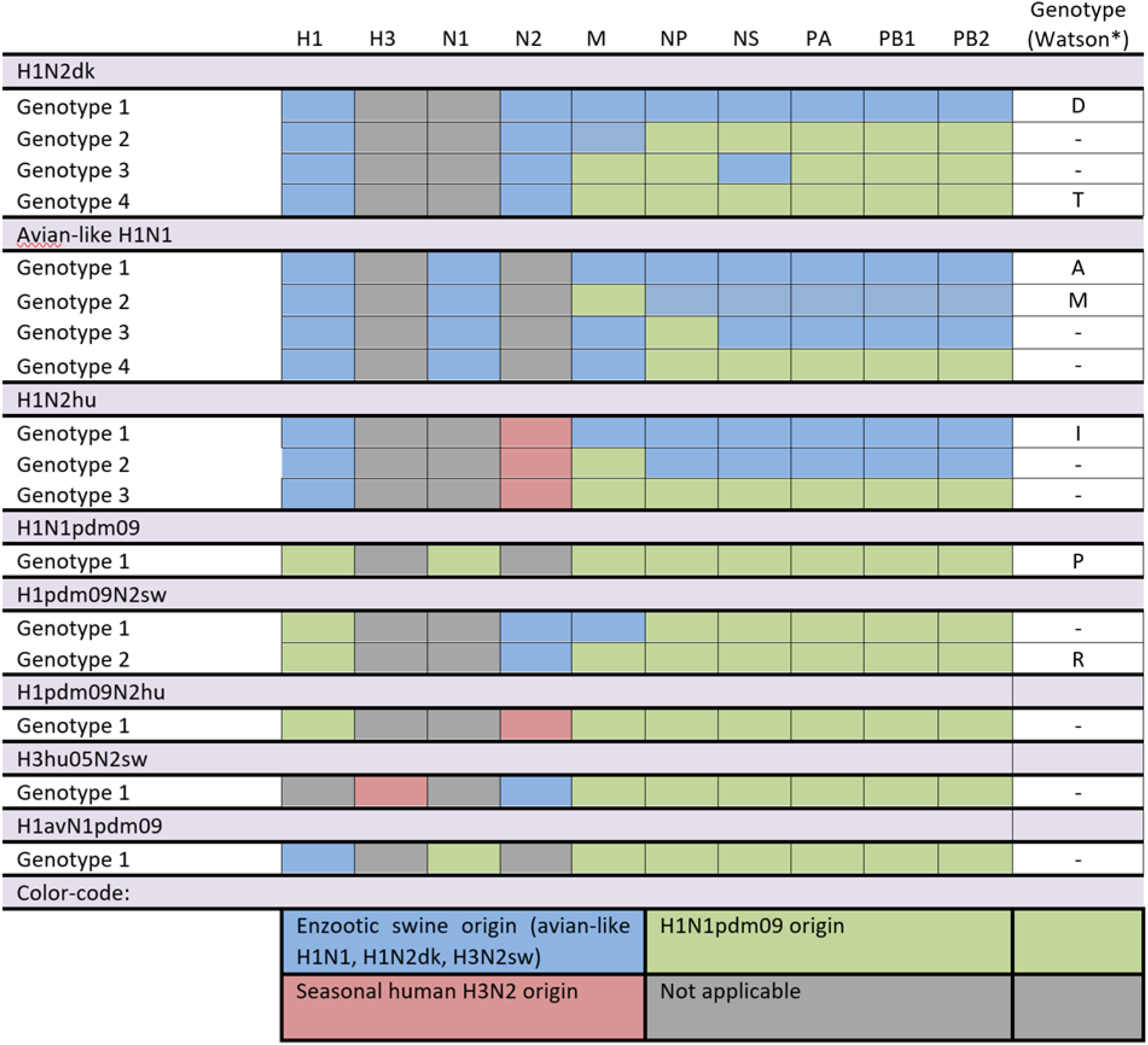
Genotypes of the different Danish swIAV isolates from 2013-2018.

All internal gene full length sequences (M, NP, NS, PA, PB1 and PB2) were subjected to individual Bayesian phylogenetic analysis, which for all gene segments revealed two main clusters; one containing sequences of avian-like swine H1N1 origin and one of H1N1pdm09 origin (Supplementary figures 8-13). Generally, there was a clear separation of the two clusters in all the phylogenetic trees. However, three and two divergent sequences were observed in-between the two main clusters in the M- and PB2 Bayesian tree, respectively (Supplementary Figure 8 and 13). The two PB2 sequences diverted due to smaller deletions, but still showed the highest sequence identity to the H1N1pdm09 subtype when performing a BLAST search. The three M sequences did not contain any deletions and the BLAST search indicated that two of the sequences (A/swine/Denmark/2013-10-1545-1p1 and A/swine/Denmark/2015-04790-1p1) shared the highest sequences identity to avian-like swine origin sequences and the third sample (A/sw/Denmark/2015-04811-10p1) shared the highest sequence identity to H1N1pdm09 origin sequences.

Full genome sequencing of the swIAV isolates obtained over the eight years, revealed that since 2013, an increasing number of the H1N2dk subtypes sequenced had acquired an internal gene cassette of H1N1pdm09 origin (Fig. 9). Similarly, though not as many samples were available, the H1avN2hu also seemed to gain internal genes of H1N1pdm09 origin over time. In contrast, the avian-like swine H1N1 subtype, roughly maintained an avian-like swine internal gene cassette, with an exception of three isolates, which contained an NP gene, an M gene and the NS, NP, PA, PB1 and PB2 genes of H1N1pdm09 origin, respectively. All other subtypes including at least one surface gene of H1N1pdm09 origin (H1N1pdm09, H1pdmN2sw, H1avN1pdm09 and H1pdmN1av) contained a complete H1N1pdm09 internal gene cassette, with the exception of one H1pdmN2sw virus (A/sw/Denmark/2013-10-1325-5p1), which had an avian-like swine M-gene. In addition, all the three full genome sequences of the H3hu05N2sw subtype contained a complete internal gene cassette of H1N1pdm09 origin (Supplementary table 2).

**Fig. 9.**
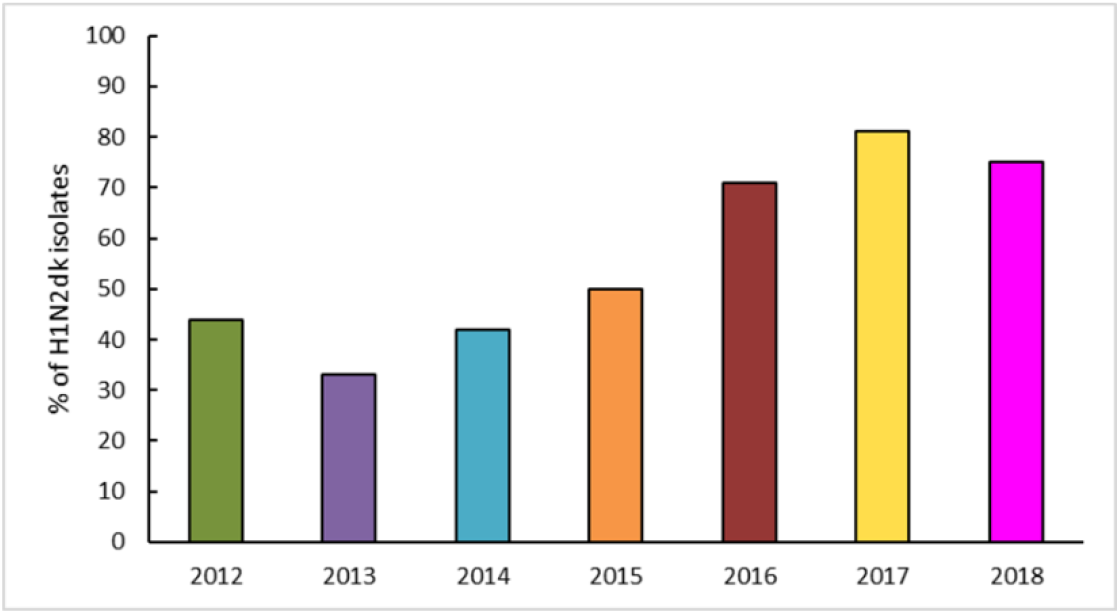
Percentage of H1N2dk isolates containing at least one gene of H1N1pdm09 origin. For the year 2011 very few sequences of H1N2dk was available and therefore the data was not included in the figure.

Previously defined important residues of the proteins encoded by the internal gene cassette was analyzed and the results are summarized in Supplementary table 3. Furthermore, comparisons of the proteins encoded by the internal gene cassette of the Sw-L and Hu-L H1pdm09Nx viruses were performed and some aa differences between the two groups were identified. However, none of the aa differences were 100 % specific to each group (Supplementary Table 4).

## Discussion

### Seasonality

IAV infections in swine has been considered a disease of late autumn and early winter^30–33^, but the results reported here reveal that while the percentage of samples testing positive for swIAV fluctuate between months, no significant differences are observed between the different seasons. This supports the recent studies describing the enzootic persistence of swIAV^34–37^, most likely as a consequence of the herd-sizes and management procedures under the current conditions of commercial swine herds. A similar lack of seasonality was found in other countries with comparable management structures^16,33,38^. Inadequate information on the severity of clinical signs were available for the Danish submissions, but a recent study from France revealed that the clinical symptoms encountered during the winter months were more severe^39^, which may explain the observed increase in the number of submissions in the autumn and winter. The increase in submissions during the autumn and winter may also be explained by the seasonal appearance of other respiratory pathogens such as mycoplasma and other bacteria^40^. Finally, some veterinarians are still considering swIAV to be a seasonal disease and are therefore not submitting samples for swIAV testing during summer. In addition, no seasonality was documented for the prevalence of H1pdm09 positive submissions, which is in accordance with a recent French study^41^. This could indicate that while H1N1pdm09 reverse-zoonosis events occurs during the human influenza season, the high level of H1N1pdm09 circulating in Danish pigs independent of the human influenza season hide the impact observed on the H1N1pdm09 occurrence during the autumn and winter months.

### Prevalent subtypes and reassortant swIAV

During the first three years of the surveillance program, the two most common influenza A virus subtypes in Danish swine were avian-like swine H1N1 and H1N2dk, which harbor the same HA gene. However, soon after the first introduction of H1N1pdm09 in January 2010, this subtype rapidly spread, and has since 2014 remained the second most prevalent subtype in Denmark. The swIAV subtype H3N2sw has almost disappeared from Denmark, in line with surveillance data obtained in some other European countries such as the UK and France^16,33^. Conversely, the H1N2dk has been steadily increasing in prevalence since 2012, and is currently the most dominating subtype in Denmark. Concurrently, the H1N2dk has gradually gained an internal gene cassette of H1N1pdm09 origin, suggesting that this gene constellation is beneficial for the virus. In general, an increase in Danish swIAV subtypes carrying an internal gene cassette of H1N1pdm09 origin was observed, which indicates that an internal gene cassette of H1N1pdm09 origin is advantageous, compared to an avian-like swine H1N1 derived internal gene cassette. The benefit of having a complete or partial internal gene cassette of H1N1pdm09 origin, could be explained by the polymerase genes having a better/increased replication efficiency^42^. In addition, certain gene combinations might enhance the transmissibility of the virus, e.g. the M-gene of H1N1pdm09 origin in combination with the A/Puerto Rico/8/34 (H1N1) strain shown increased transmissibility in the guinea pig model^43^. Interestingly, based on the phylogenetic trees, it seems that the internal gene cassette might have an influence on the evolution of the surface genes, as several clusters among the H1av, N2sw and N2hu sequences correlates with the origins of the internal gene cassette. However, further studies are needed to investigate how the different gene segment can influence each other, but it might be related to the specific reassortment event forming a common ancestor for the cluster. Finally, the replacement of the avian-like swine internal gene cassette with an H1N1pdm09 internal gene cassette, could enhance the zoonotic potential, as proposed for the American H3N2v^44^ and the British H1N2r^45^ subtypes, which have resulted in several human infections. Therefore, the pandemic potential of swIAV harboring gene segments of H1N1pdm09 origin should be a future research focus.

Six novel reassortant swIAV subtypes and a total of 17 genotypes were identified during the eight-year surveillance period. These findings underline the importance of having a national swIAV surveillance program, which acts as an early warning system both for the swine industry and for the human health sector, ensuring that novel subtypes and variants escaping current vaccines can be quickly identified. The H3hu05N2sw subtype is a perfect example hereof, as it is a triple-reassortant swIAV including gene segments from IAV of enzootic swIAV origin, H1N1pdm09 origin and human seasonal IAV origin^26^. Surprisingly, this subtype has only been sporadically detected during the last five years. A possible dissemination of this subtype among Danish swine herds would probably have devastating consequences, because there is no population immunity towards the human seasonal H3hu05 ^26^. This indicates that other factors than pre-existing immunity towards the HA protein are important for the spread of novel swIAV subtypes and strains. Indeed, the most successful virus in Denmark during the last seven years has been the H1N2dk, despite that there has been a high level of population immunity towards the HA protein of this subtype since the 90’s. Combined with the findings that the internal cassette of H1N1pdm09 origin seems to benefit viral competitiveness, we might need to change our perception that pre-existing immunity to HA is the main driver of evolution to focus also on the impact of the internal genes. Two other cases of human seasonal IAV spillover into the swine population were observed during the surveillance, including the H1avN2hu and H1pdmN2hu subtypes. Both subtypes contain the NA gene of a human seasonal IAV circulating in the 90’ties^29^. The continued circulation of the H1pdmN2hu subtype in swine is worrying from a zoonotic perspective, because all eight gene segments of this virus originates from viruses known to be able to replicate in- and spread between humans. The circulation of the H1avN2hu subtype is even more worrying, since there is no immunity against the HA protein of this subtype in the human population. The H1avN2hu has gradually gained the internal cassette of H1N1pdm09 origin, meaning that some of these viruses contain seven out of eight gene segments, which have been found in human IAV strains and thereby may lead to increased zoonotic potential. Therefore, it is important to monitor the occurrence of these subtypes in the future – both in pigs and in humans. Another group of reassortant swIAV, that potentially pose a problem for the swine herds, are those mixing the surface genes of enzootic swIAV and H1N1pdm09 subtypes. These novel reassortants includes H1pdmN2sw, H1pdmN1av and H1avN1pdm. Swine herds experiencing infections with one of these three subtypes could potentially have a reduced effect of vaccination, as no available vaccines currently include both the H1N1pdm09 subtype and the enzootic swIAV subtypes. Thereby, these farms might need to apply two vaccines to reach an optimal immunity to the circulating herd strain.

### Genetic and antigenic drift

Another important aspect of swIAV evolution is the genetic drift, mainly affecting the two surface genes (HA and NA)^46,47^. Especially the avian-like swine hemagglutinin protein (H1av) seem to have undergone extensive genetic and antigenic drift, as a great sequence diversity was revealed. It was evident that the evolution of the H1av gene did not evolve in one specific direction over time, but rather evolved in many different directions, resulting in a vast number of different H1av clusters and variants. In a recent study^37^, we found that the evolution of the H1av in a single herd followed a pectinate pattern mirroring the pattern seen globally for the human seasonal influenza strains, which contradict the general perception that swine IAV is not prone to selection driven by preexisting immunity like in humans. In the present study, we assessed the H1av evolution over time in the Danish pig population as a whole and over several years and failed to confirm this pectinate pattern. Thus, it seem that swIAV evolution at the single herd level is identical to the pattern seen in the global human population, but when the swIAV evolution is evaluated on a national or global scale, this pectinate pattern is disrupted. The reason for this difference is probably that the human population, due to the extensive global interactions, can be regarded as a single “epidemiological unit”, whereas swine herds, due to a high level of external biosecurity and limited exchange of live animals between herds, represents a vast variety of closed “epidemiological units”, which each have a specific- and probably pectinate-pattern of evolution. In other words, the global evolution of swIAV is characterized by a vast number of different local clusters of viruses that on the herd level evolve similar to human seasonal influenza viruses. This in turn results in very complex phylogenetic trees with a lot of clusters and subclusters, which disrupt the pectinate structure. Still, despite the lack of a clear pectinate like evolution, the H1av variants had clearly undergone positive selection on specific codons located in antigenic sites, which is known to alter the binding of neutralizing antibodies^48–50^. This further confirm our previous findings, that the herd immunity leads to significant antigenic drift in the globular head of the HA protein, as seen for human seasonal IAV^51^. Furthermore, the finding of similar residues undergoing positive selection between different herds, indicates that some residues in the HA protein are of particular importance for swIAV evolution. Finally, the substitution rate estimated for H1av was similar to that documented in previous studies^52–55^, but was lower than the substitution rate estimated for H1av in a single herd over time^37^. This emphasize that one should differentiate when comparing evolutionary results based on data obtained from a single herd or data obtained through extensive surveillance programs.

The H1pdm09 sequences analyzed in this study, revealed the existence of two groups of sequences. One group of H1pdm09 sequences forming a well-defined cluster only containing sequences derived from swine (Sw-L sequences) and another group of more diverse swine derived H1pdm09 sequences (Hu-L sequences) that were scattered among human seasonal H1pdm09 sequences. In general, relatively long branches separated the Danish Hu-L H1pdm09 swine sequences and the closest human sequences. However, a few of the Hu-L sequences from Danish swine had a high level of identity to viruses isolated from humans during the corresponding human influenza season, indicating a very recent “spill-over” from humans to pigs. Indeed, all the Danish Hu-L viruses probably represents reverse zoonotic events, where the H1N1pdm09 virus was transmitted from humans to swine, and has started to evolve in pigs and by that has drifted away from the human “seed” virus. The other group of H1pdmNx viruses found in Danish swine, formed a clearly defined cluster (Sw-L cluster) that was different from the human seasonal H1N1pdm09 sequences. Interestingly, this cluster also contained 11 viruses isolated from swine in Germany and one from Italy. This is not surprising, since Denmark has an annual export of more than 10 million weaned pigs to mainly Eastern Europe and Germany, which are not tested for swIAV prior to export. In contrast, all swine adapted H1N1pdm09 viruses (SwD) found in France during recent years^41^ formed another unique cluster that were only distantly related to the “Danish-German” Sw-L cluster, confirming that the evolution of swIAV follows different evolutionary traits in populations that are not epidemiologically connected.

We estimated that the diversion into the Sw-L clusters occurred around 2011, approximately one year after the first H1N1pdm09 virus was detected in Danish swine. This indicates that the virus needed little time to become established in pigs, which is supported by the finding that this subtype constituted 21 % of the IAV positive samples already in 2011. Comparison of the aa sequences between the two Danish H1pdm09 clusters (Sw-L and Hu-L) revealed 20 aa differences and four of these aa positions were shared between the Danish Sw-L cluster and the French swine divergent cluster (SwD)^41^, indicating that these residues are important for adaption of this virus to swine. The fact that several of the 20 aa differences were present in the RBS emphasize the probable relation to host-adaption.

In summary, the presented data strongly indicate that the human seasonal H1N1pdm09 viruses still are capable of infecting swine, despite more than ten years of adaption to humans, but it is unclear if the swine adapted viruses of the Sw-L cluster also have retained its capability to infect humans. Studies to investigate this in the ferret model are ongoing.

In comparison to the H1av sequences, the H1pdm09 sequences exhibited a lower level of sequence diversity, probably because the H1pdmNx has circulated in Danish swine for significant shorter time than the H1avNx strains. In contrast, the substitution rate and the positive selection on the RBS and antigenic sites were comparable to that of the H1av sequences when the evolutionary analysis were performed on the Danish H1pdm09 Hu-L sequences separately, whereas there was no evidence of positive selection on the H1pdm09 Sw-L sequences. Moreover, the substitution rate and the temporal signal were higher for the Hu-L sequences compared to the Sw-L sequences. Nevertheless, as described above, 20 aa residues were identified that differed between all or most of the H1pdm09 sequences of the Sw-L cluster and Hu-L sequences. Thirteen of these changes were situated in the RBS or antigenic sites, indicating that the two groups of viruses had been under separate selective pressure. Additionally, the Sw-L H1pdm09 proteins seemed to have gained changes enhancing O-linked glycosylation in connection to position 150 located in the RBS. The general epidemiological differences between these two groups of viruses is that the Sw-L group has circulated among pigs since 2011, whereas the Hu-L group probably represents multiple introductions from humans in different seasons and by that have had less time to adapt to pigs. Thus, the evolutionary rate calculated for the Hu-L sequences may actually reflect evolution that happened in humans prior to the jump into pigs. This notion is supported by no specific swine and human residues being identified when comparing all swine and human derived H1pdm09 sequences. Another explanation could be that the specific differences observed in the H1pdm09 protein sequences of the Sw-L group of viruses reflect adaption to swine, which probably mainly took place during the first passages among pigs, whereafter the sequences mainly experience negative selection as seen for the Sw-L group of viruses that diverted into a separate clusters around 2011. There is a lack of published data on the molecular adaptations that take place during these reverse zoonotic events of influenza A virus and therefore the hypotheses described above remain speculative. From a zoonotic point of view, it is worrying that the H1N1pdm09 viruses seem to evolve in different directions in pigs and humans, especially if the swine adapted Sw-L viruses retain their capacity to infect humans. Thirteen of the 20 aa residues, that differed between the human and swine adapted viruses, were situated in important antigenic sites or the RBS, and 7/20 aa residues were present in all Sw-L H1pdm09 sequences and were absent in the Hu-L and human seasonal H1pdm09 sequences. However, antigenic cartography performed on H1N1pdm09 viruses collected in France, showed a high degree of cross-protection between the swine adapted and the human seasonal-like H1N1pdm09 viruses isolated in 2014-16^41^. Nonetheless, it should be taken into consideration that the Sw-L cluster of our study showed several changes different form the French swine divergent cluster (SwD) and therefore the antigen cartography should be repeated on the Danish H1pdmNx viruses of the Sw-L cluster. Overall, the monitoring of the antigenic evolution of H1pdmNx swine adapted viruses should be prioritized in the future, to ensure early detection of emerging virus with altered antigenicity. This is highly important, as decreased cross protection between these two clusters would be detrimental if the swine adapted virus jumps back into humans. Similarly having an IAV monitoring of personal in affected herds should be considered.

### Specific host and virulence markers

In summary, the HA proteins of the H1pdm09 viruses seem to be better adapted to elicit a strong receptor binding to the α2.6-linked sialic acid receptor compared to the H1av HA proteins. This may reflect that the H1pdm09 HA are descendants of the H1N1 “Spanish flu” strain^18^ and by that have circulated in mammals for at least 100 years, whereas the H1av HA protein was first detected in a mammal (pig) in the eighties’^56^. In turn, these results could also explain why very few cases of zoonotic infection involving H1av have been registered^57,58^. The fact that more Danish Hu-L sequences had “D” at position 225 support the assumption that these H1pdmNx viruses are indeed more similar to human seasonal-like H1Npdm09 viruses compared to the viruses of the Sw-L cluster and also indicate that the G225D transition may be more important in humans than in swine. The comparison of swine H1pdm09 sequences and human seasonal H1pdm09 aa sequences revealed that the residue at position 273 might be a potential marker important for distinguishing between swine and human H1pdm09 sequences. However, this residue was not 100 % unique to one of the two groups of sequences, and more studies should be performed to identify specific swine and human markers of the H1pdmNx subtypes. In addition, the eight aa residues defined to differ between avian IAVs and H1N1pdm09 origin viruses in the NP, PB1, PB2 and PA gene segments^59^ were consistent with the residues observed in the two clusters (avian-like swine and H1N1pdm09) of the NP, PB1, PB2 and PA genes segments of this study. This suggests that these residues are indeed specific for H1pdmNx swIAV.

The recently identified residues 48Q, 98K and 99K of the Eurasian avian-like swIAV NP protein conferring MxA resistance^60^, was documented in the majority (81%) of the Danish NP protein of avian-like swine origin. MxA resistance is essential for zoonotic and pandemic potential of avian and swine IAV^60,61^, and there is therefore a potential increased risk of zoonotic transmission in the Danish herds, where circulation of swIAV strains carrying these three mutations is present.

As for the aa changes observed between the Sw-L and Hu-L sequences in the internal proteins the T76A change in the PB2 protein has been linked an elevated interferon response^62^. Furthermore, the M283I aa change in the PB2 protein has previously been linked to decreased virulence of avian H5 IAV^63,64^ and the N456S aa change has, on the other hand, been linked to human adaptations of the H3N2 subtype^65^. For the PB1 protein, the M317I aa change has been identified in a H2N2 after multiple passaging in chicken eggs to create a temperature sensitive IAV strain^66^. Finally, the C241Y in the PA protein has been linked to mammalian adaptions of avian H5N1 viruses^67^. In summary, several of the aa changes observed between the Sw-L and Hu-L internal proteins have previously been described to have an influence on the virulence, replication efficiency or host-response/adaptation, thereby emphasizing that these changes could be important in the adaption of H1pdm09Nx viruses to swine. However, this needs to be investigated further.

### Importance of swIAV surveillance programs

The results generated in connection with the passive surveillance program of swIAV performed in Denmark from 2011-2018, highlights the importance of such a program. The surveillance was essential in identifying novel subtypes and variants that circulate among Danish swine, and the knowledge supports veterinarians and farmers daily in selecting the most compatible swIAV vaccine and understanding the swIAV transmission dynamics in the herd. Moreover, novel subtypes not covered by the current available vaccines were identified, thereby avoiding unnecessary use of vaccines and encouraging medical companies to prioritize vaccine-updates. These vaccine updates are not only encouraged by identifying novel subtypes, but also by documenting the level of antigenic drift, which previously has been shown to affect the level of cross-protection between strains of the same lineage^68^. However, HI-tests, antigenic cartography, virus neutralization assays and finally controlled animal experiments should be performed on a range of different strains within each lineage to investigate the consequence of the genetic drift on the cross-protection. The number of submissions for swIAV diagnostics increased the last years of the surveillance, indicating that the program is useful for farmers and veterinarians. Moreover, an increase in the number of submissions positive for swIAV may indicate that swIAV infections represent an increasing problem in Danish swine herds or that there is increased focus on swIAV as an important pathogen in the herds. Finally, the ability of the program in identifying novel subtypes and variants that might have an increased zoonotic potential is vital from a human health perspective, as it can function as an early warning system for future human IAV pandemics.

## Materials and Methods

### Samples

Samples, including lung tissues, nasal swabs and oral fluids, originating from swine herds experiencing clinical signs of acute respiratory disease, were submitted for routine diagnostic examinations at the Danish National Veterinary Institute by veterinary practitioners from 2011 until 2018. The submissions included 1 to 5 samples (yearly average: 2-2.9) from each herd.

### RNA isolation

Total RNA was extracted from lung tissue, nasal swab samples or cell cultured virus isolates by RNeasy Mini Kit (QIAGEN, Denmark) automated on the QIAcube (QIAGEN) according to the instructions from the supplier. The samples were prepared for extraction as follows; 200 μl nasal swab sample or virus isolate were mixed with 400 μl RLT-buffer containing β–mercaptoethanol, whereas 30 mg of lung tissue was homogenized in 600 μl RLT-buffer containing β–mercaptoethanol for 30 sec at 15 Hz using the TissueLyser II (Qiagen).

Oral fluid samples were prepared by homogenization of 200 μl sample (30 sec at 15 Hz) in a Tissuelyser II (Qiagen) followed by centrifugation (2 min at 10.000 rpm). Total RNA was extracted from 140 μl of the prepared oral fluid sample using the QIAamp Viral RNA Mini Kit (Qiagen) automated on the QIAcube (Qiagen) according to the instructions from the supplier.

The total RNA from all sample types was eluted in 60μl RNase-free water and stored at −80 °C until further analysis. Positive and negative controls were included in all extractions.

### Detection of swIAV

The presence of swIAV was detected by an in-house modified version of a real time RT-PCR assay detecting the M gene^69^. The assay was performed in a total reaction volume of 25 μl using the RNA Ultrasense One-Step Quantitative RT-PCR System (Invitrogen), 3 μl of extracted RNA, 300 nM forward primer (RimF), 600 nM 5’-labeled reverse primer (MaR-FAM), 400 nM 3’-labeled probe (MaProbe). Details of the primers and probes are listed in Table 1. All reactions were analyzed on the Rotor-GeneQ machine (Qiagen) using the following PCR conditions: [50 °C 30 min; 95 °C 2 min; 45 cycles of 95 °C 15 sec, 55 °C 15 sec (acquiring using 470 nm as source and 660 nm as detector), 72 °C 20 sec; 95 °C 15 sec; Melt curve analysis by ramping from 50 °C to 99 °C, wait for 90 sec on pre-melt conditioning at first step, rising by 1 °C each step and wait for 5 sec before acquiring]. A positive and negative control were included in all runs.

### Test for the HA gene of H1N1pdm09 origin by specific real time PCR

All swIAV positive samples were tested for the presence of the HA gene of H1N1pdm09 origin (H1pdm09) by an in-house real time RT-PCR assay detecting specifically the HA gene of the pandemic virus (Table 1). All reactions were analyzed in a Rotor-GeneQ machine (Qiagen) using the following PCR conditions: [45 °C for 10 min; 95 °C for 10 min; 45 cycles of 95 °C for 15sec; 55 °C for 20 sec; 72 °C for 30 sec]. In 2018, an additional assay targeting the H1pdm09 was implemented to increase the sensitivity of the H1pdm09 screening (Table 1). The two H1pdm09 assays were run as a multiplex on the Rotor-GeneQ machine (Qiagen) using the following PCR conditions: [45 °C, 20 min; 95 °C, 15 min; 45 cycles: 94 °C, 30 sec; 55 °C, 20 sec; 60 °C, 20 sec]. A positive and negative control were included in all runs.

### Subtyping

From 2011-2014, the swIAV positive samples were subtyped using Sanger sequencing of the HA and NA genes according to a previously described PCR protocol^70^. The PCR products were purified using the High Pure PCR product Purification Kit (Roche, Denmark). Subsequently the purified PCR products were sent for sequencing at LGC Genomics (Berlin, Germany) with primers comprised of the “pQE” part of the PCR primers (Table 1).

From 2015-2017, samples were subtyped using a multiplex real time RT-PCR assay strategy. Two multiplex reactions including primers and probes for H3hu, N1pdm, H1av, N2hu or H3sw, H1pdm, N1sw, N2sw+hu, respectively^34^ (Table 1) were analyzed on the Rotor-GeneQ machine (Qiagen) using the following PCR conditions: 50 °C for 20 min; 95 °C for 15 min; 40 cycles of 94 °C for 60 sec, 60 °C for 90 sec. In 2018, subtyping of swIAV positive samples were in addition performed on the Fluidigm PCR platform (AH diagnostics, United States) according to a previously published protocol^71^. All runs on the Rotor-GeneQ and the Fluidigm included positive controls representing all the possible subtypes targeted by the different assays along with a negative control.

### Virus isolation

Virus was isolated from selected swIAV positive clinical specimens by inoculation of Madin Darby Canine Kidney (MDCK) cells following standard cell culture procedures. In short, 150 mg lung tissue was homogenized in 1.5 ml MEM (Invitrogen Carlsbad, CA, USA) containing 1000 units/ml Penicillin and 1 mg/ml Streptomycin. Sterile filtrated inoculums were prepared in viral growth medium (MEM 1x, L-Glutamin 2 mM, Non-essential amino acids 1x, 100 units/ml Penicillin, 100 μg/ml Streptomycin and TPCK-treated trypsin 2 μg/ml) using either 10 % lung tissue homogenate or 20 % nasal swab or oral fluid sample. The inoculum was added to 70 % confluent MDCK cells for 45 minutes at 37 °C and 5 % CO2 followed by the addition of fresh viral growth medium after wash of the inoculated cells. After 3 days, the cell culture supernatant was harvested and tested for influenza A virus by real time RT-PCR.

### Full genome sequencing

From 2013-2017 full genome sequencing was performed on cell culture-propagated influenza virus samples, which had been subjected to full-length PCR amplification of all eight gene segments with in-house designed primers (Table 1) using SuperScript III OneStep RT-PCR System with Platinum Taq High Fidelity. The PCR conditions were as follow for each gene segment: HA: 55°C, 30 min, 94 °C, 2 min, 4x (94 °C, 30 sec – 55 °C, 30 sec - 68 °C, 180 sec), 41x (94 °C, 30 sec – 68 °C, 210 sec) and 68°C, 10 min. NA: 54 °C, 30 min, 94 °C, 2 min, 4x (94 °C, 30 sec – 58 °C, 30 sec - 68 °C, 180 sec), 41x (94 °C, 30 sec – 68 °C, 210 sec) and 68°C, 10 min. M: 50°C, 30 min, 94 °C, 2 min, 41x (94 °C, 30 sec – 56°C, 30 sec - 68°C, 90 sec) and 68°C, 10 min. Nucleoprotein (NP): 58 °C, 30 min, 94 °C, 2 min, 4x (94 °C, 30 sec – 54 °C, 30 sec - 68 °C, 180 sec) and 41x (94 °C, 30 sec – 68 °C, 210 sec) and 68°C, 10 min. Nonstructural protein (NS): 58°C, 30 min, 94 °C, 2 min, 41x (94 °C, 30 sec – 55°C, 30 sec, 68°C, 90 sec) and 68°C, 10 min. Polymerase basic protein 1 (PB1) and polymerase acidic protein (PA): 52 °C, 30 min, 94 °C, 2 min, 4x (94 °C, 30 sec – 52 °C, 30 sec - 68 °C, 180 sec), 41x (94 °C, 30 sec – 68 °C, 210 sec) and 68°C, 10 min. Polymerase basic protein 2 (PB2): 55 °C, 30 min, 94 °C, 2 min, 4x (94 °C, 30 sec – 52 °C, 30 sec - 68 °C, 180 sec), 41x (94 °C, 30 sec – 68 °C, 210 sec) and 68°C, 10 min. Purified PCR products for all gene segments were pooled in equimolar quantity to a final amount of 1 μg and used for next generation sequencing (NGS) on the Ion Torrent PGM™ sequencer. The NGS, including library preparation, was carried out at the Multi-Assay Core facility located at the Technical University of Denmark. In 2018, full genome sequencing were performed on cell culture propagated virus samples using universal influenza primers^72^ (Table 1). Library preparation and NGS on the Illumina MiSeq platform were conducted at the Statens Serum Institut, Denmark.

### Sequence analysis

Data obtained from Sanger sequencing and subsequent analyses of the consensus sequences were performed using CLC Main Workbench version 7.6.2-20.0.3 (CLC bio A/S, Aarhus, Denmark). Alignments of each gene segment were created using the MUSCLE algorithm^73^. Phylogenetic trees were constructed using a distance-based method with the Neighbor Joining algorithm and bootstrap analysis with 1000 replicates. The results were verified by using Maximum Likelihood Phylogeny. Sequences obtained by NGS were assembled using the features “de novo assembly” and “map read to references” using 22 reference sequences representing the different lineages of each gene segment in CLC Genomics Workbench 4.6.1-8.0.2 (CLC bio A/S). The subtype and lineage of each sample and gene segment were determined based on MUSCLE alignments, subsequent neighbor joining phylogenetic trees, and the function “BLAST against NCBI”. Moreover, sequence alignments of each lineage of the two surface gene segments (H1pdm09, H1av, N1pdm, N1av, N2hu and N2sw) were analyzed for the average nucleotide diversity (pi) using author’s own software. For more detailed phylogenetic analysis, Bayesian trees of each gene segment (internal genes) and lineage (H1pdm09, H1av, N1pdm, N1av, N2hu and N2sw) were constructed using the program MrBayes with the following settings; nst=mixed and rates=invgamma. The trees were run for 10.000.000 generations and a sample frequency of 500^74^. An additional alignment and Bayesian tree was constructed for the H1pdm09 gene, including all available European swine H1N1pdm09 sequences from NCBI GenBank and GISAID and all Danish human H1N1pdm09 sequences available for the years 2009-2018 in GISAID together with a selection of human H1N1pdm09 sequences from other countries. For visualization, the number of sequences were subsequently reduced excluding sequences with 100 % nucleotide sequence identity. A list of all the reference sequences used can be found in Supplementary table 1. Convergence of the Bayesian analysis was checked using Tracer version 1.7.1^75^, and the results visualized in Figtree version 1.4.4^76^.

In addition to the Bayesian phylogenetic analyses, strict molecular clock trees were constructed for the surface gene segments of the lineages; H1pdm09, H1av, N1pdm, N1av, N2hu and N2sw to determine the temporal evolution and the substitution rate. However, before the trees were constructed, all sequences were investigated for the presence of a temporal signal (i.e., whether nucleotide changes accumulate roughly proportionally to elapsed time) using the program TempEst^77^ and evaluating the correlation coefficient. Subsequently, the alignments of each lineage were analyzed in the program BEAST2 version 2.5.2, where the model settings were as previously described^68^. Briefly, the HKY substitution model with gamma-distributed rates over sites was chosen along with a strict clock model including tip dates. The outcome of the analysis was visualized in Figtree version 1.4.4^76^ and convergence checked in Tracer version 1.7.1^75^.

The surface gene segments of the different lineages; H1pdm09, H1av, N1pdm, N1av, N2hu and N2sw were investigated for the presence of positive selection using the CODEML program of the PAML package as previously described^48^. Briefly, this was done by comparing the fits of CODEML’s substitution models M1a and M2a (NSsites = 1 and 2). M1a includes two categories of codons – some under negative selection (dN/dS ratio < 1) and some codons where mutations are neutral (dN/dS ratio = 1). Model M2a includes three categories of codons – the same two as M1a plus an additional category of codons under positive selection (dN/dS ratio > 1). If M2a fits a dataset significantly better than M1a, then there is evidence of positive selection in some codons (and the identity of these codons is also determined during model fitting). The average dN/dS ratio (global ω ratio) of the surface gene segments of the different lineages; H1pdm09, H1av, N1pdm, N1av, N2hu and N2sw were also estimated using CODEML with the setting NSsites = 0.

All nucleotide sequences of each gene segment were translated into amino acid (aa), and MUSCLE^73^ alignments were created using CLC Main Workbench 20.0.3 (CLC bio A/S, Aarhus, Denmark). Subsequently, the alignments were manually examined to determine the presence of previously described aa differences and residues. Specifically, for the HA proteins these included residues unique to the H1pdmN2sw subtype^27^ and residues linked to receptor binding^78,79^. Moreover, the five previously defined antigenic sites Sa, Sb, Ca1, Ca2 and Cb of the H1 subtype^49,80,81^ and the receptor-binding site (RBS)^82^ were annotated to the H1av and H1pdm09 proteins and investigated for divergence and correlations to codons with increased dN/dS ratios. For the NA protein residues encoding neuraminidase inhibitor resistance were investigated^83^. All PB2 proteins were examined for specific residues encoding virulence^84^, pathogenicity ^85^ and host adaptation^86^. The eight residues of the NP, PB1, PB2 and PA proteins proposed to differ between avian viruses and viruses of the H1N1pdm09 subtype^59^, were also investigated. Finally the three residues of the NP protein recently found to confer MxA resistance^60^ were examined. The two groups of the H1pdm09 proteins were examined for differences in the number and location of N-linked and O-linked glycosylation sites using the NetNGlyc 1.0^87^ and NetOGlyc 4.0^88^ servers from DTU Bioinformatics, Denmark.

### Statistics

Results of the screening for swIAV and H1pdm09 in each submission were analyzed in Microsoft Excel 2016 version 16.0.4993.1001 and GraphPad^89^. The proportions of swIAV positive, swIAV negative and the proportion of H1pdm09 positive submissions compared to total number of swIAV positive submissions were calculated for each month. The monthly average percentage of swIAV positive submissions and the proportion of H1pdm09 positive submissions were calculated based on the results obtained from each month during the eight years, and the differences in the pairwise percentages and proportions of swIAV and H1pdm09 submission were investigated using a student’s t-test and a Fisher’s exact test in GraphPad^89^. To determine differences between the prevalence of a specific aa residue at a given position a chi-squared test in GraphPad^89^ was utilized. P-values below 0.05 were considered statistically significant.

## Acknowledgements

The authors would like to acknowledge all the Danish herds that submitted samples for the surveillance. Moreover, we acknowledge the authors, originating and submitting laboratories of the sequences that we obtained from GISAID’s EpiFlu™ Database (www.gisaid.org) and NCBI GenBank (www.ncbi.nlm.nih.gov).

## Competing interests

The authors declare no conflict of interest.

## Funding

The farmers or the medical company IDT Biologika GmbH paid the initial screening for the presence of swIAV in a submission, while the Danish Veterinary and Food Administration paid the remaining analyses. In addition, the work presented in this article was supported by Novo Nordisk Foundation (FluZooMark – grant # NNF19OC0056326)

## Supplementary tables

**Supplementary table 1.**
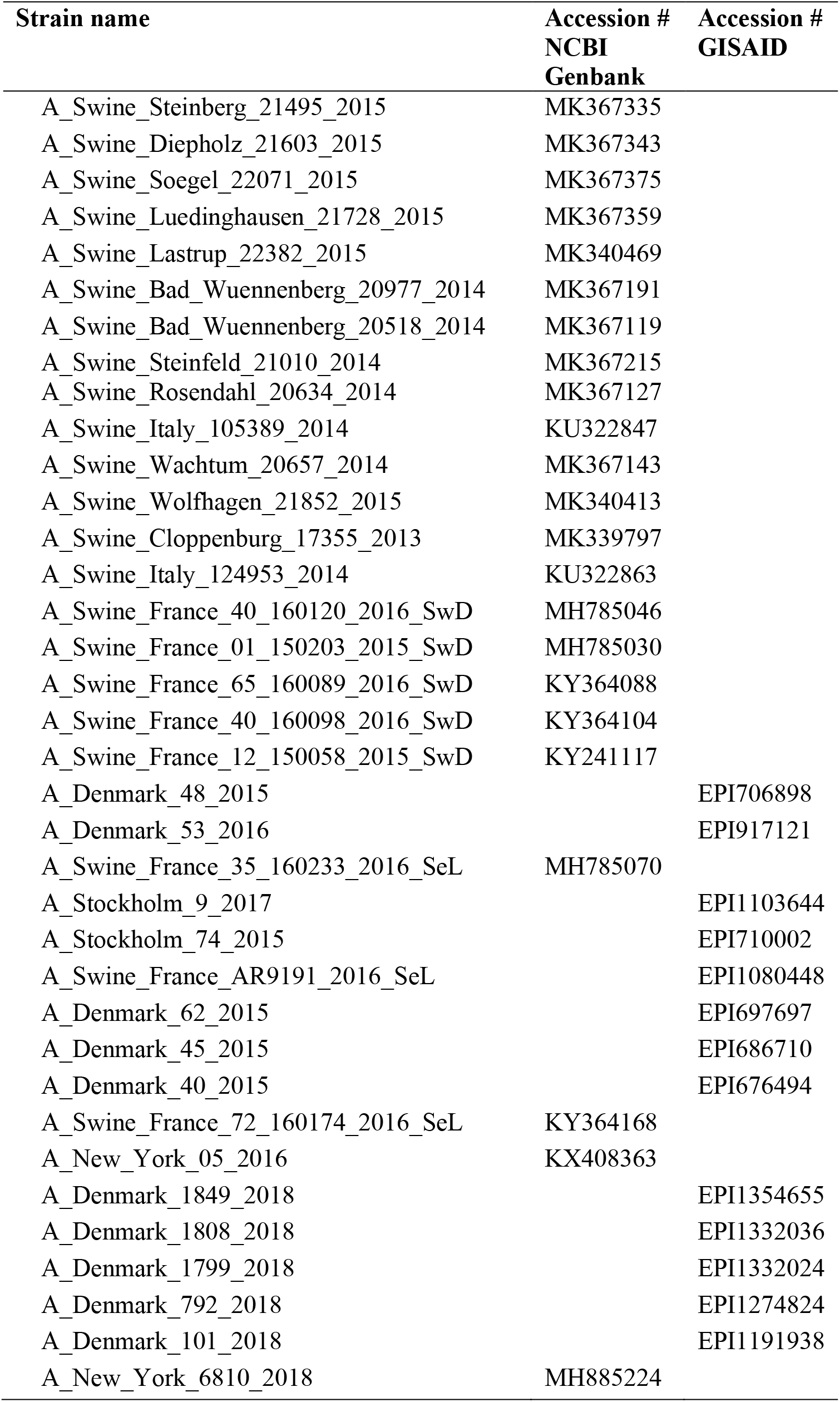
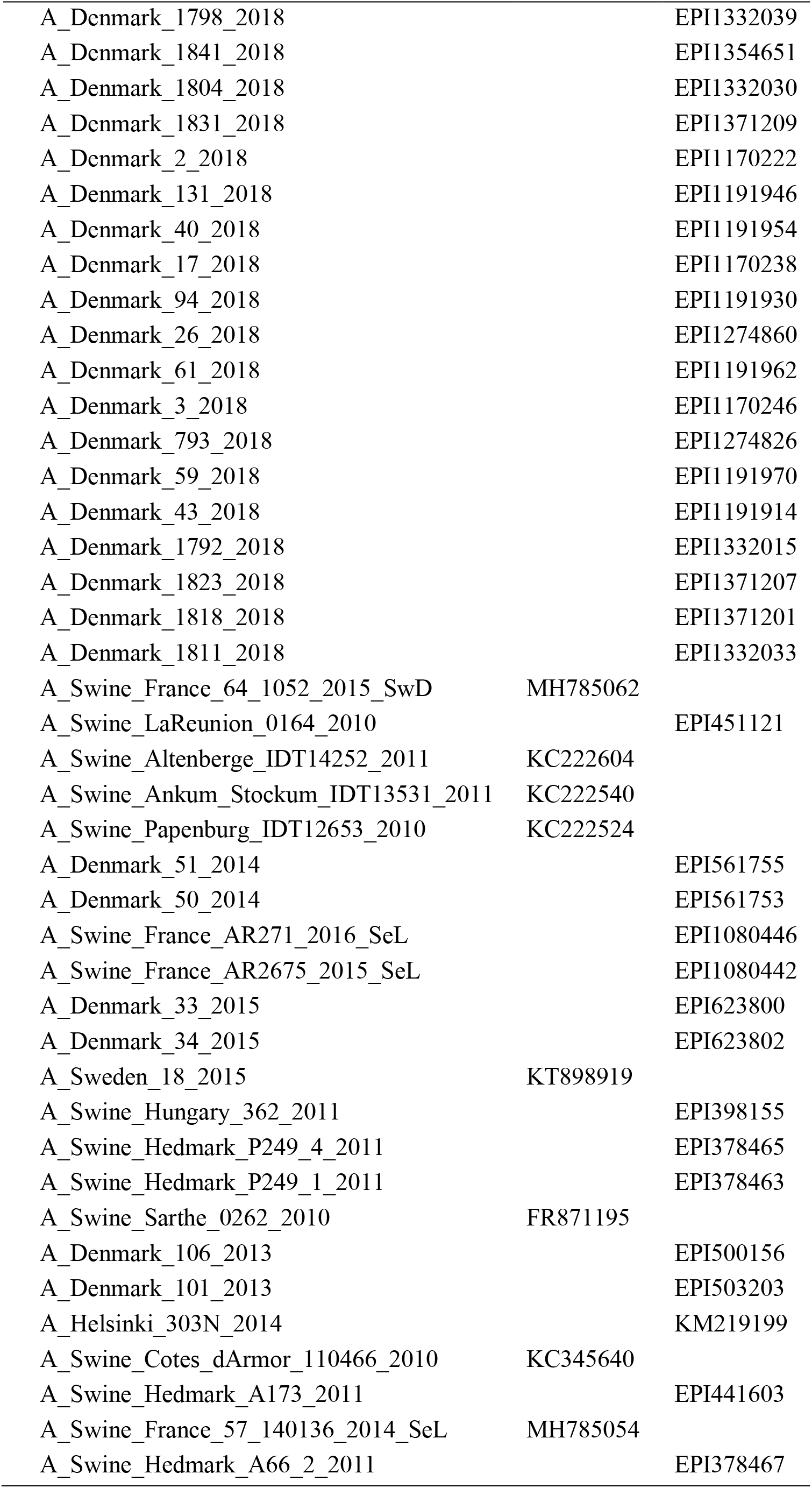
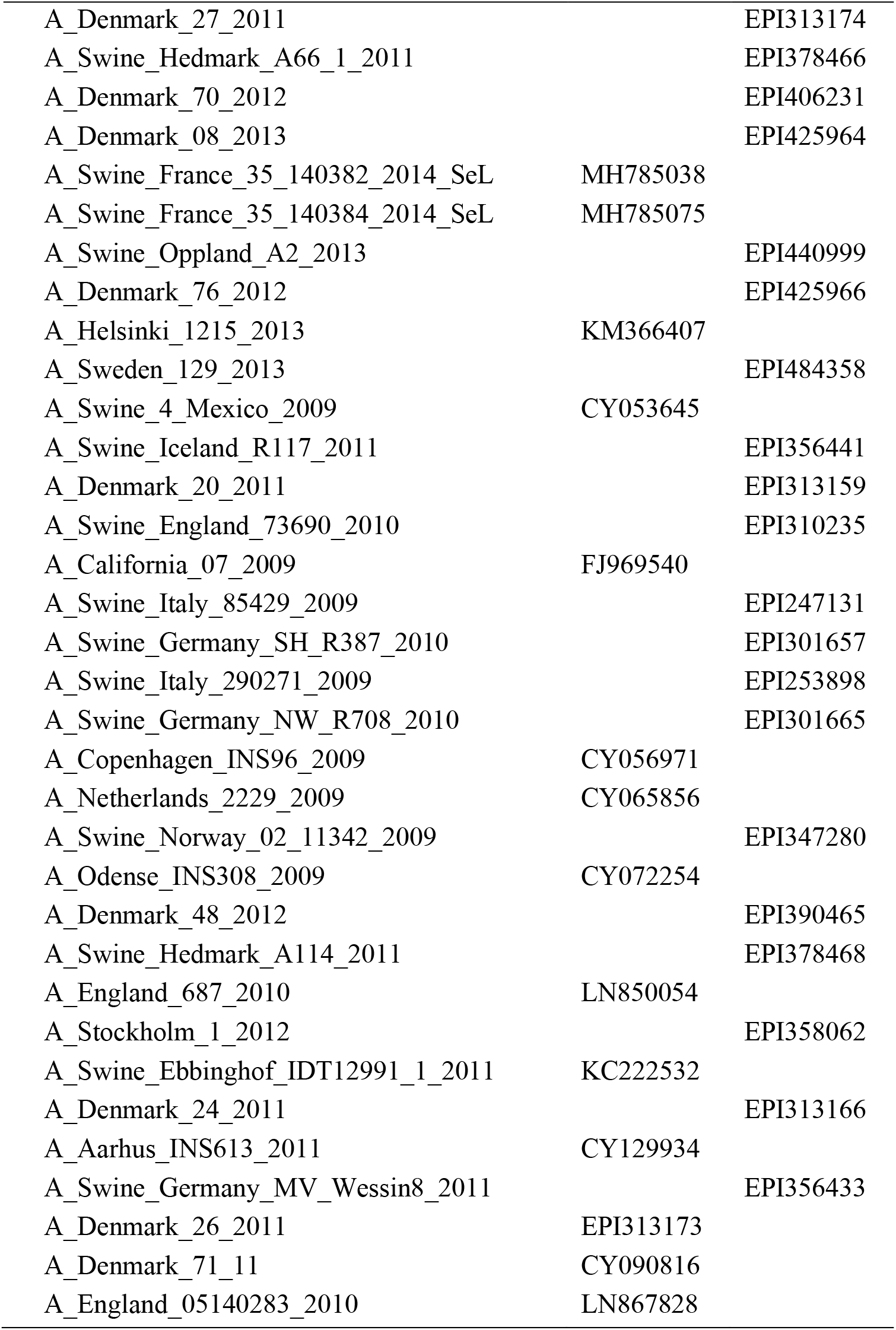
list of H1pdm09 reference sequences included in Fig 7.

**Supplementary table 2.**
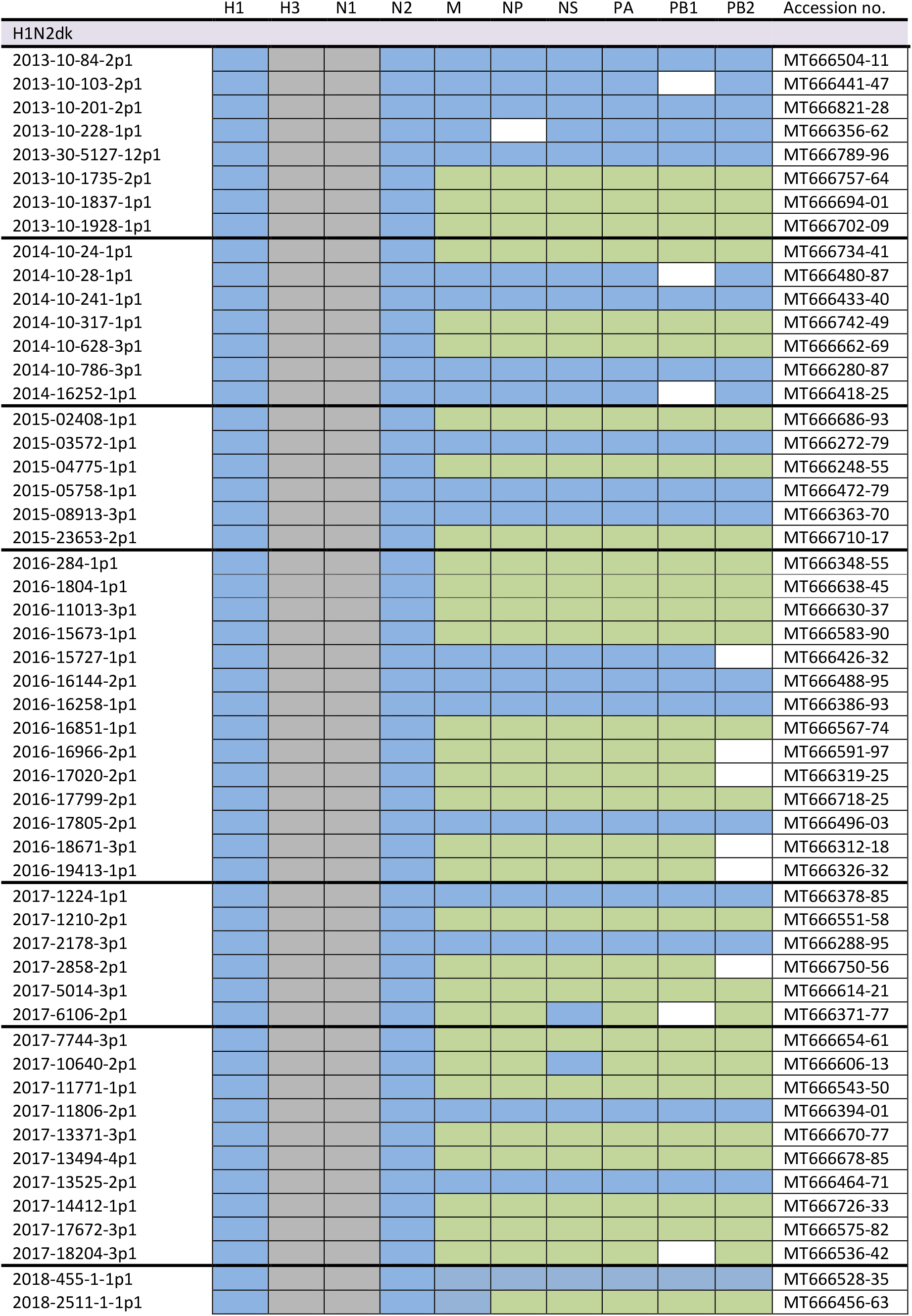
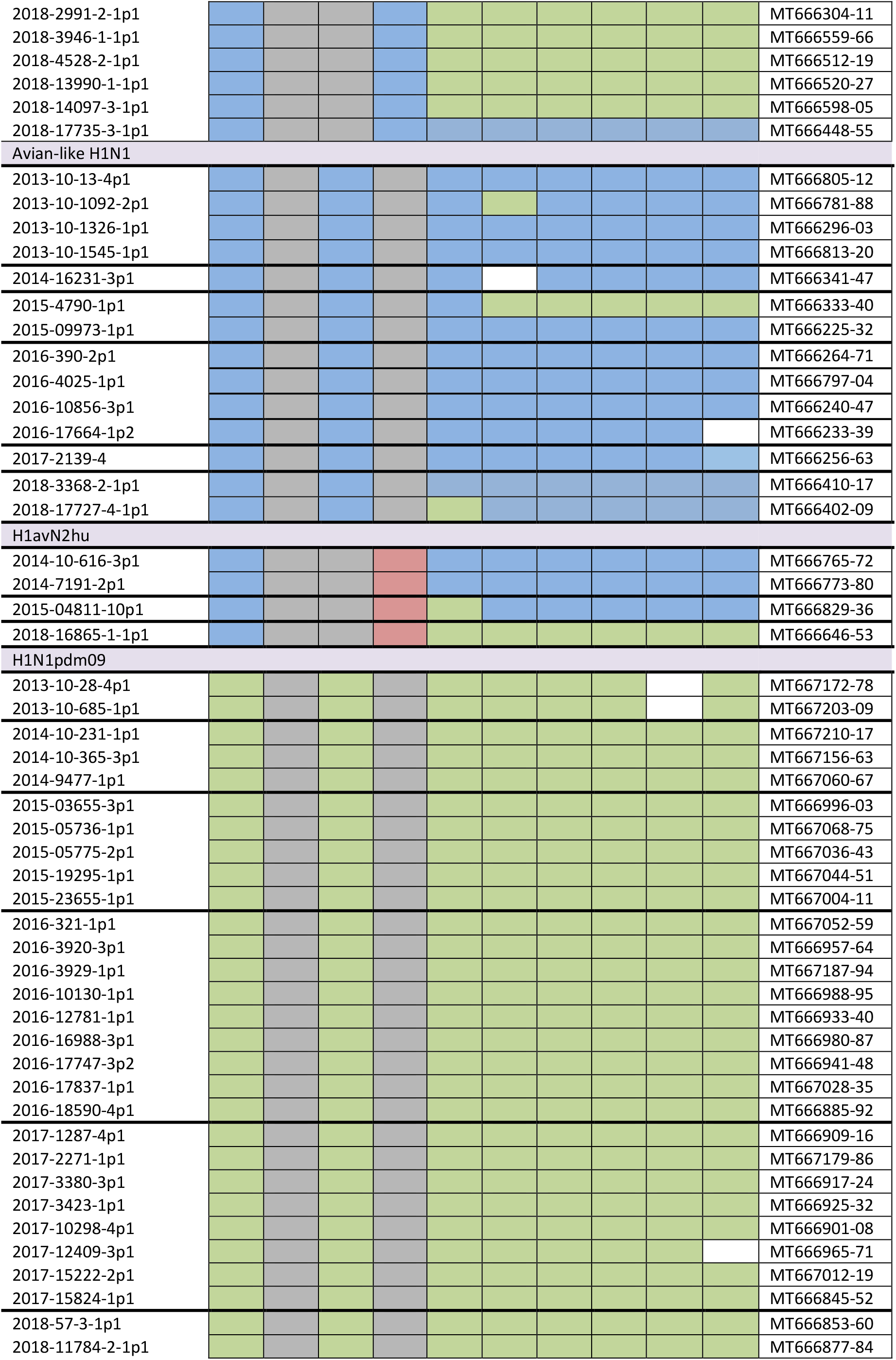
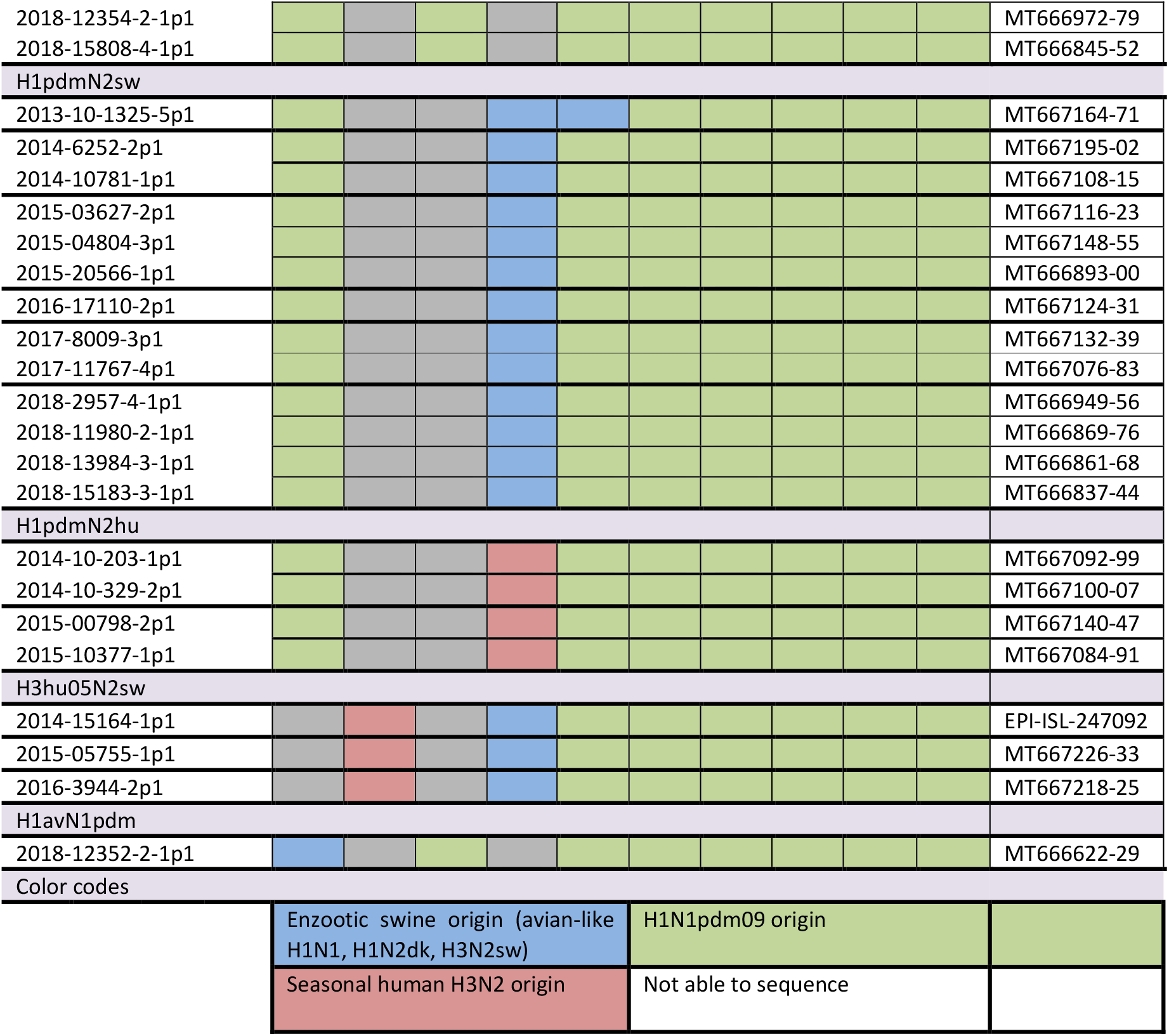
the genotype of all full genome sequenced samples

**Supplementary table 3.** residues examined for specific mutations involving host adaptation, virulence, pathogenicity and dominating residues differing between avian-like and H1N1pdm09 origin viruses.

| Protein and residue/aa change | Function | H1av-origin | H1N1pdm09 origin |
| --- | --- | --- | --- |
| **PB2** |  |  |  |
| T271A | Dominant residue differ between avian- and H1N1pdm09 viruses<sup>59</sup> | 35/36 had T<br>1/36 had I | 85/85 had A |
| E627K | Increased viral replication and virulence<sup>84</sup> | 0/78 | 0/48 |
| D701N | Enhance viral replication and pathogenicity of Eurasian avian like swIAV<sup>85</sup> | 36/78 | 0/48 |
| K702R | Host-adaption; at this residue avian influenza carries lysine while most mammalian IAV carry an arginine<sup>86</sup> | 0/78 | 0/48 |
| **PB1** |  |  |  |
| V336I | Dominant residue differ between avian- and H1N1pdm09 viruses<sup>59</sup> | 34/37 had V<br>3/37 had I | 87/87 had I |
| **PA** |  |  |  |
| K356R | As above<sup>59</sup> | 38/38 had K | 88/91 had R |
| S409N | As above<sup>59</sup> | 5/38 had S<br>33/38 had N | 90/91 had N<br>1/91 had S |
| **HA** |  |  |  |
| T200A (H3 numbering) | Increased receptor binding affinity to the α2.6-linked sialic acid for H1N1pdm09 viruses<sup>79</sup> | 1/78 had A | 48/48 had A |
| E190D (H3 numbering) | change in receptor specificity of avian IAV from α2.3-linked sialic acid to α2.6-linked sialic acid<sup>78</sup> | 73/78<br>5/78 had V or S | 40/48<br>8/48 had S or G (all Hu-L) |
| G225D (H3 numbering) | As above<sup>78</sup> | 6/78<br>63/78 had E<br>9/78 had T/K/N | 18/48 (14/18 was Hu-L)<br>23/48 had N<br>1/48 had G |
| K159, G172E, I183V, S200P, S202N, D204S, V338I and V66I | Related to the H1pdmN2sw subtype in Germany<sup>27</sup> |  | Not specific for H1pdmN2sw. However, I173V and V466I were present in 3/12 and 1/12 of the Danish H1pdmN2sw viruses |
| **NP** |  |  |  |
| K48Q | Confer MxA resistance in combination with R98K and R99K for the NP protein of | 35/35 had Q | 92/92 had K |

|  | avian-like swine origin<sup>60</sup> |  |  |
| --- | --- | --- | --- |
| E53D | Confer MxA resistance for the NP protein of H1N1pdm09 origin<sup>61</sup> | 35/35 had E | 10/92 had D (8/10 were H1pdmNx viruses belonging to the Hu-L cluster) |
| R98K | Confer MxA resistance in combination with K48Q and R99K for the NP protein of avian-like swine origin<sup>60</sup> | 33/35 had K | 92/92 had R |
| R99K | Confer MxA resistance in combination with K48Q and R98K for the NP protein of avian-like swine origin<sup>60</sup> | 31/35 had K | 90/92 had R |
| R100I/V | Confer MxA resistance for the NP protein of H1N1pdm09 origin<sup>61</sup> and dominant residue differ between avian- and H1N1pdm09 viruses<sup>59</sup> | 35/35 had R | 84/92 had I<br>3/92 had M<br>3/92 had V<br>2/92 T/L |
| F313V | Confer MxA resistance for the NP protein of H1N1pdm09 origin<sup>61</sup> | 35/35 had F | 91/92 had V |
| V33I | Dominant residue differ between avian- and H1N1pdm09 viruses<sup>59</sup> | 8/35 had V<br>27/35 had I | 91/92 had I<br>1/92 had V |
| R100V | As above<sup>59</sup> |  |  |
| R305K | As above<sup>59</sup> | 35/35 had R | 91/92 had K<br>1/92 had R |
| Q357K | As above<sup>59</sup> | 35/35 had Q | 92/92 had K |

Aa positions are numbered according to the first methionine if nothing else is indicated.

**Supplementary table 4.**
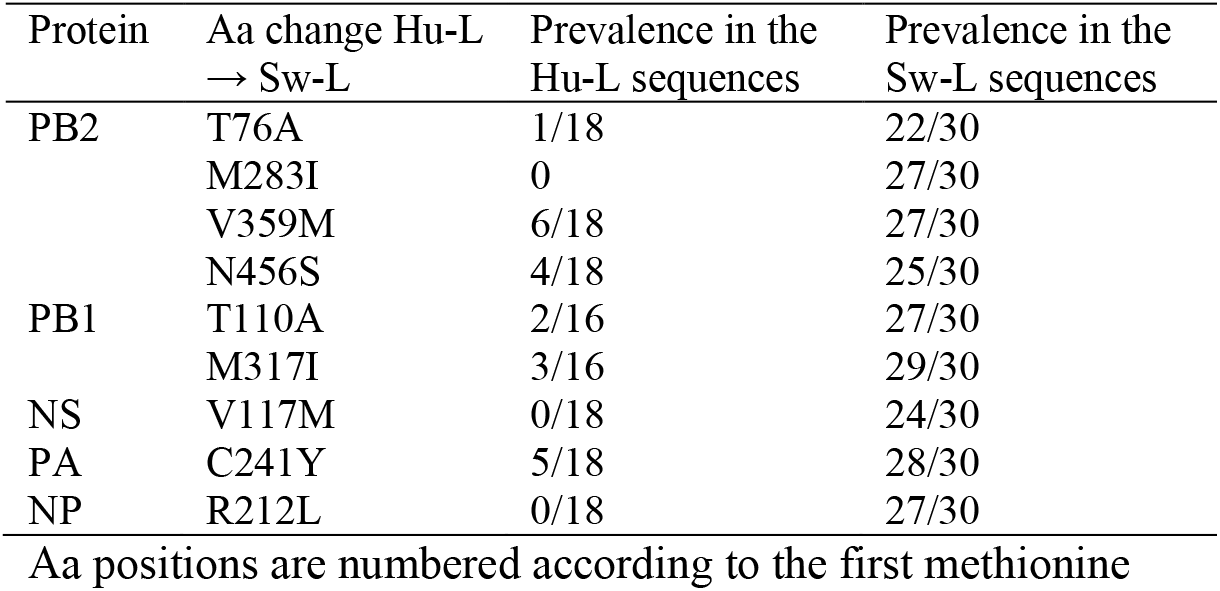
Amino acid differences in the internal proteins of the Hu-L and Sw-L H1pmd sequences

## Supplementary figures

**Supplementary figure 1.**
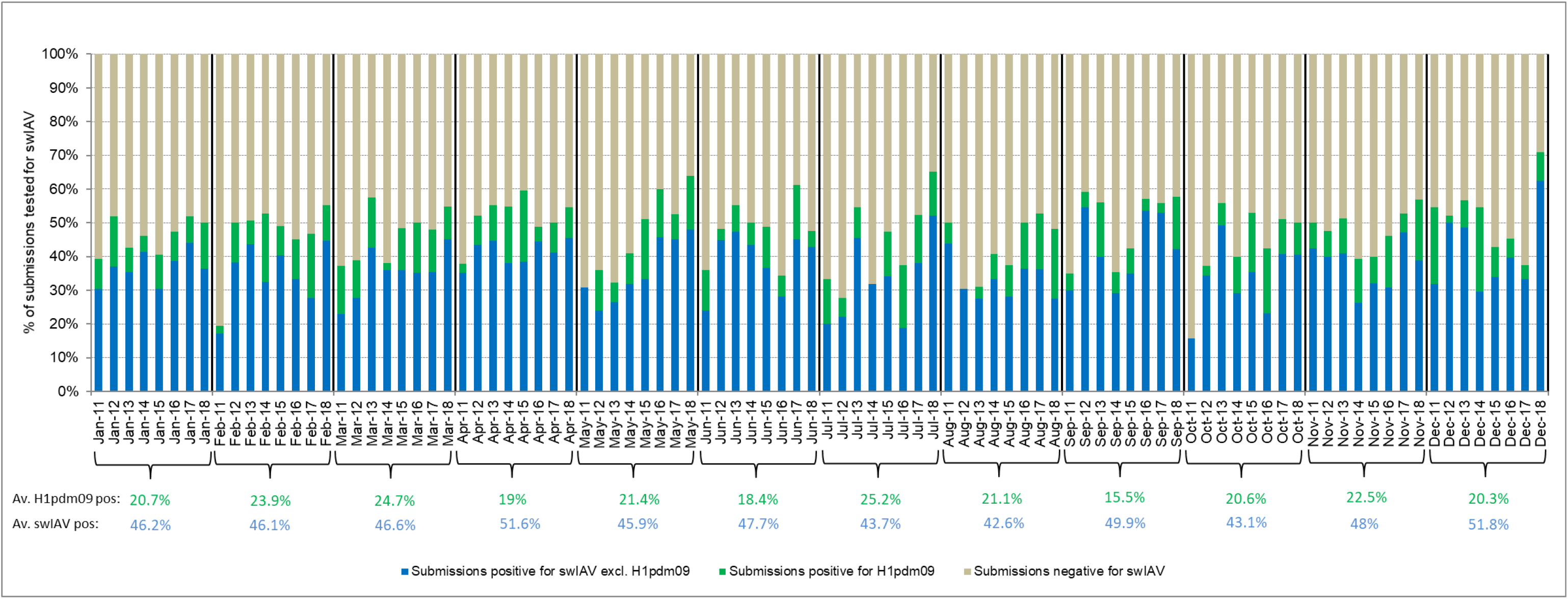
Monthly distribution of swIAV submissions 2011-2018. The average percentage of swIAV positive and H1pdm09 positive over the eight year surveillance period is indicated below each representative month in blue and green, respectively.

**Supplementary Figure 2.**
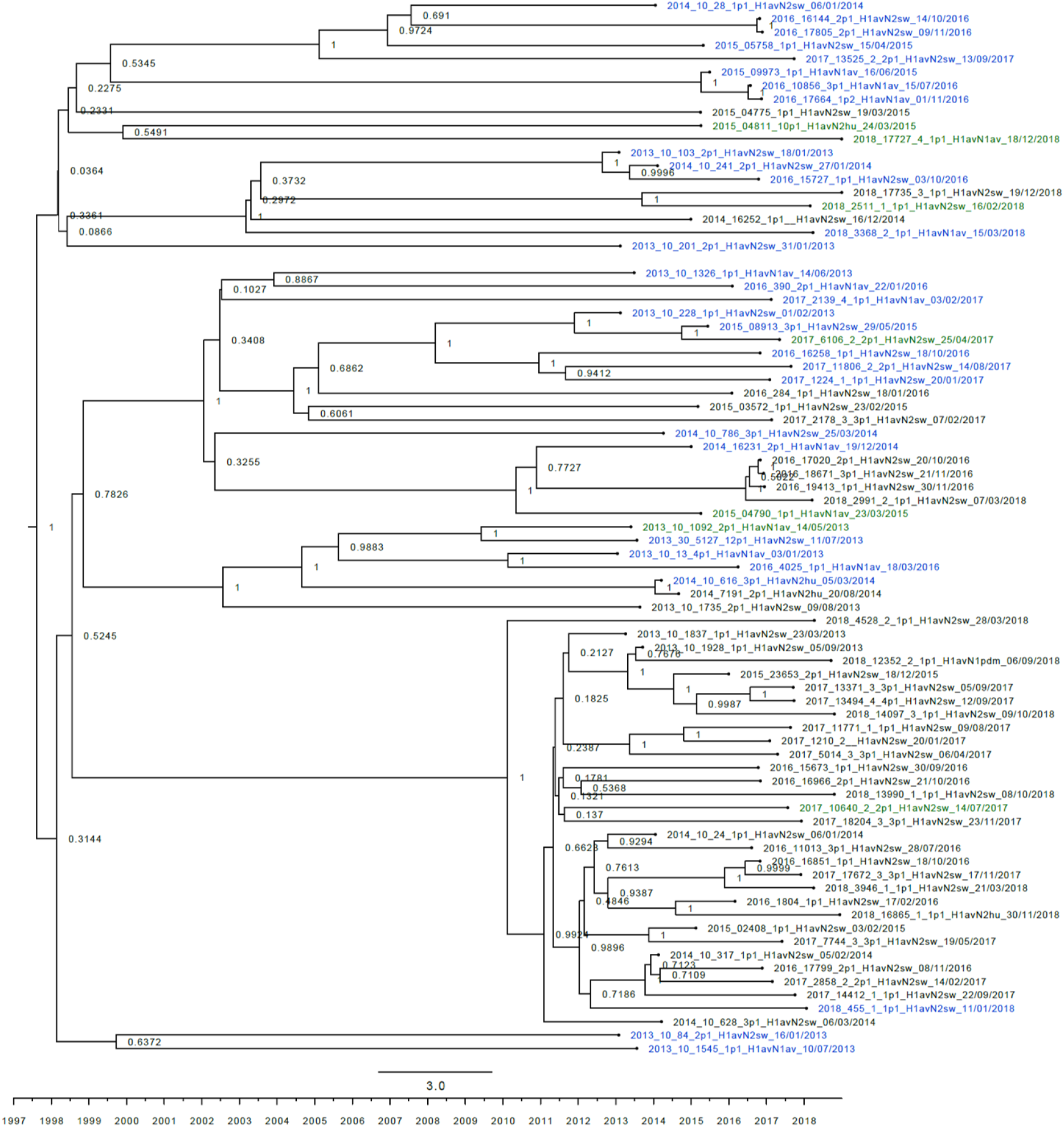
strict molecular clock tree of the H1av nucleotide sequences The x-axis indicates the time in years and each tick indicates half a year. A blue taxon indicates that the sample carried an internal gene cassette of avian origin, whereas a green taxon indicates that the sample carried a partial internal gene cassette of H1N1pdm09 origin. A black taxon indicates that the sample carried an internal gene cassette of H1N1pdm09 origin. The x-axis represents time in years.

**Supplementary Figure 3.**
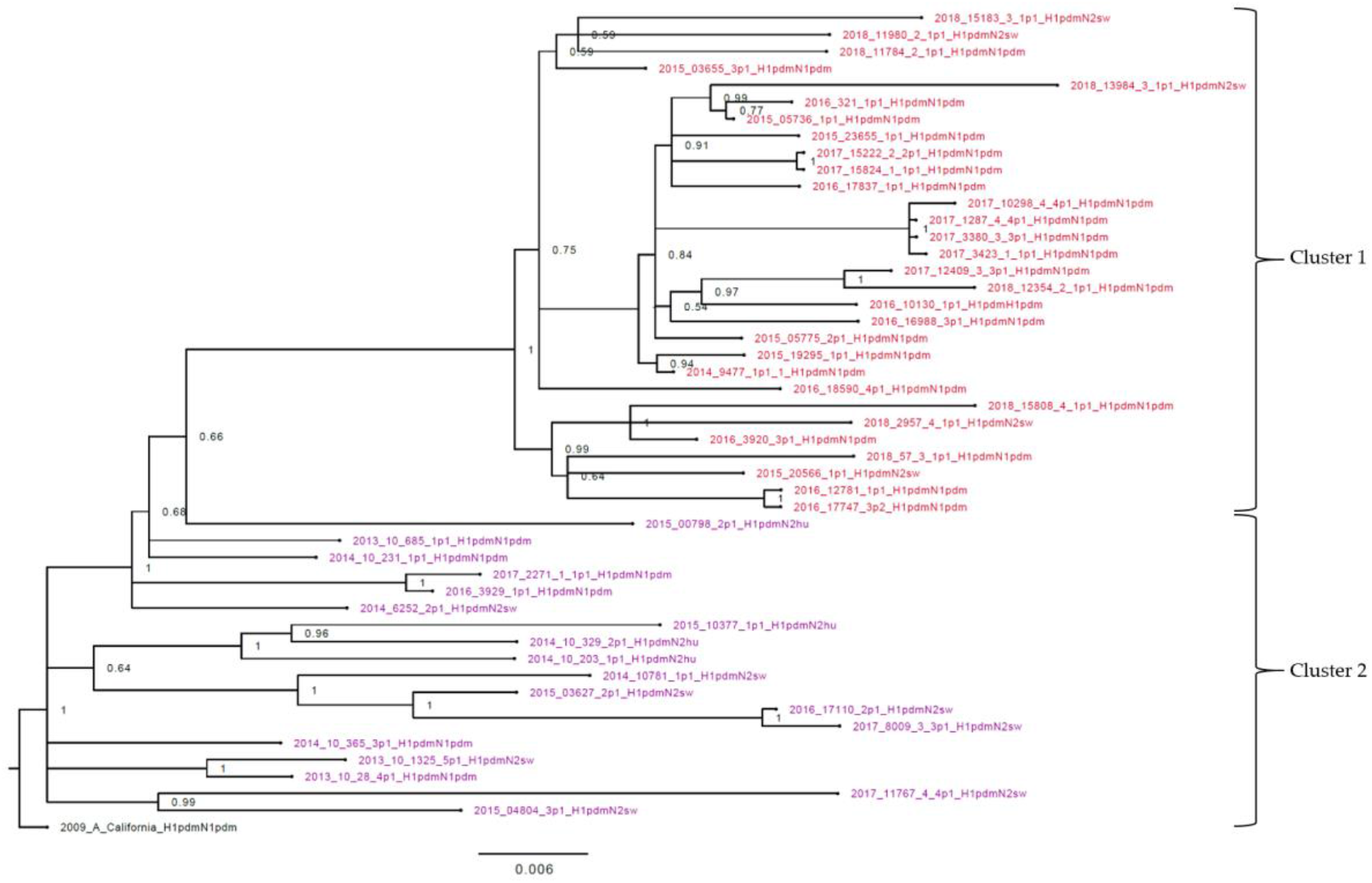
Bayesian phylogenetic tree of the H1pdm nucleotide sequences Node labels indicate posterior probabilities. “2009_A_California” is the outgroup. A red taxon indicate that the sequence is part of Cluster 1 and a purple taxon indicates that the sequence is part Cluster 2.

**Supplementary Figure 4.**
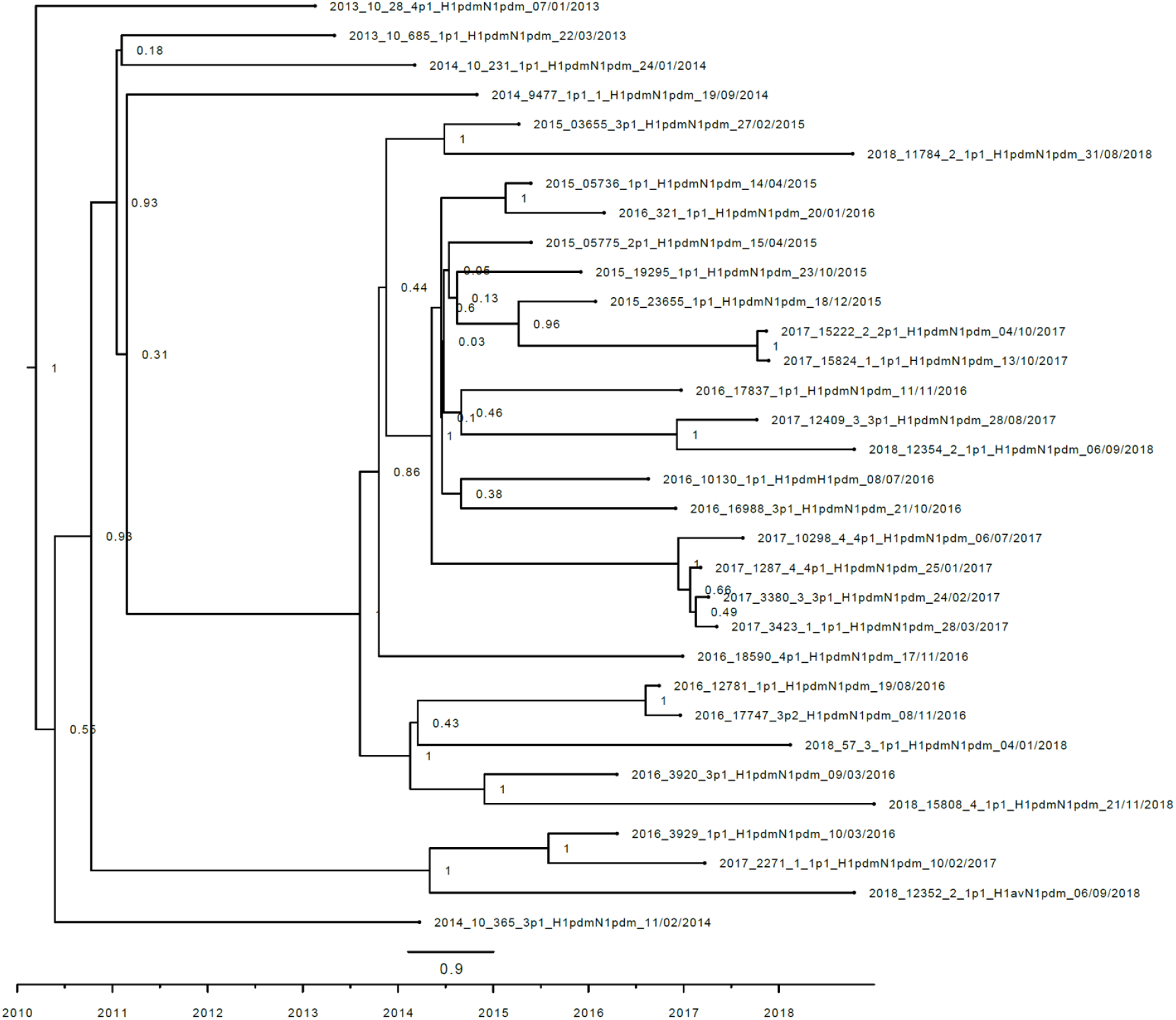
Strict molecular clock tree of the N1pdm sequences The x-axis indicates the time in years and each tick indicates half a year. A black taxon indicates that the sample carried an internal gene cassette of H1N1pdm09 origin.

**Supplementary figure 5.**
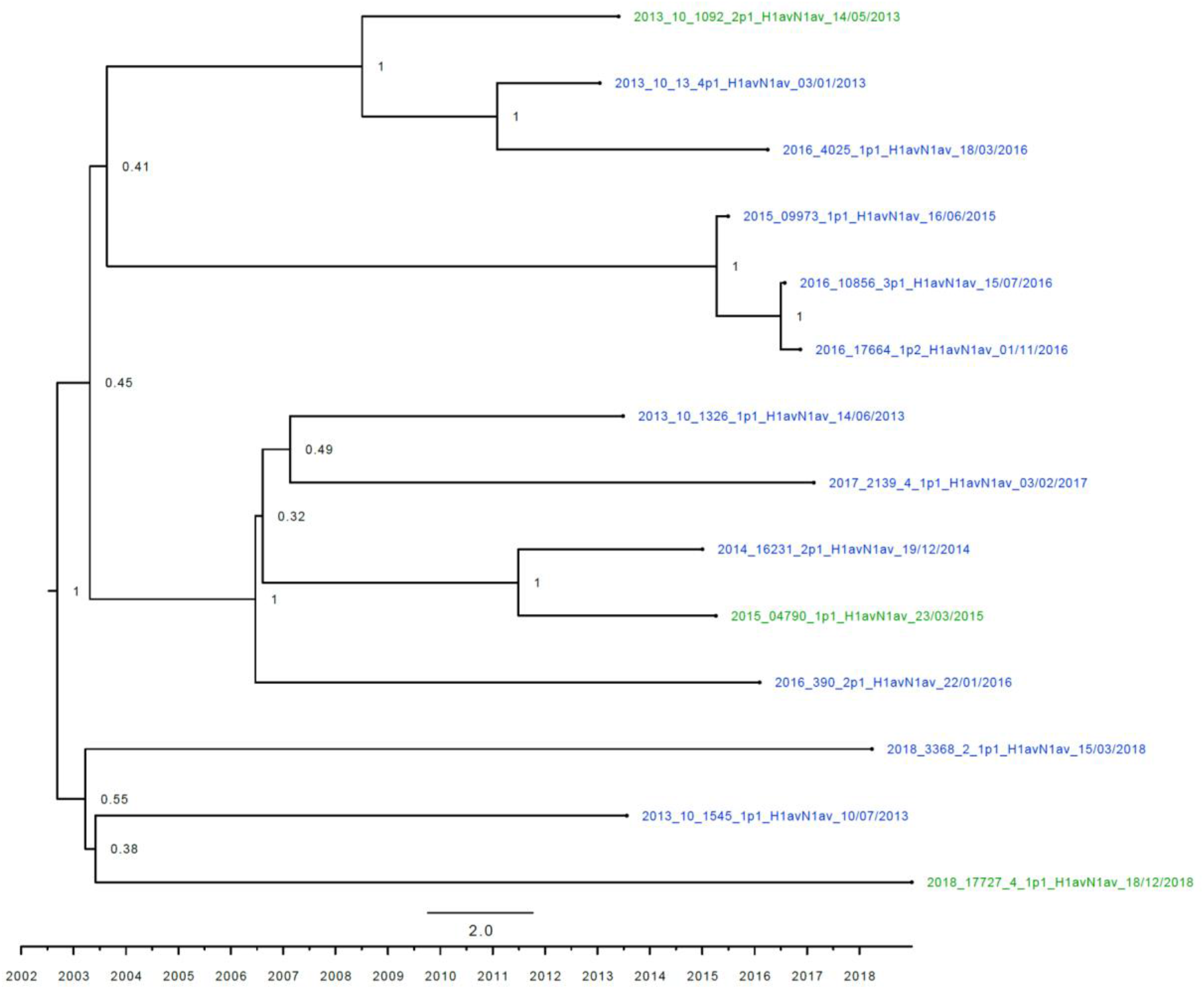
Strict molecular clock tree of the N1av sequences The x-axis indicates the time in years and each tick indicates half a year. A blue taxon indicates that the sample carried an internal gene cassette of avian origin, whereas a green taxon indicates that the sample carried a partial internal gene cassette of H1N1pdm09 origin.

**Supplementary figure 6.**
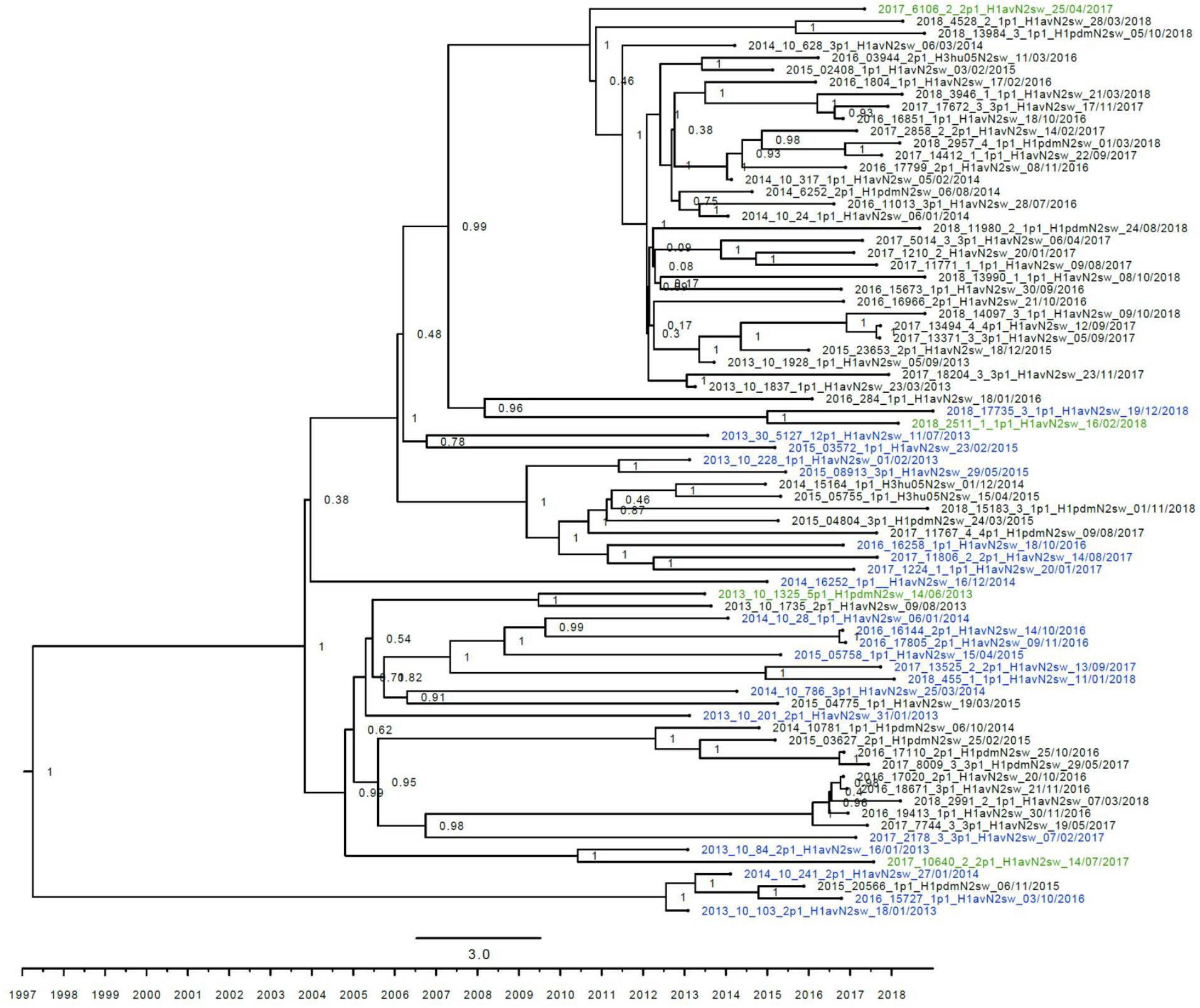
Strict molecular clock tree of the N2sw sequences The x-axis indicates the time in years and each tick indicates half a year. A blue taxon indicates that the sample carried an internal gene cassette of avian origin, whereas a green taxon indicates that the sample carried a partial internal gene cassette of H1N1pdm09 origin. A black taxon indicates that the sample carried an internal gene cassette of H1N1pdm09 origin.

**Supplementary figure 7.**
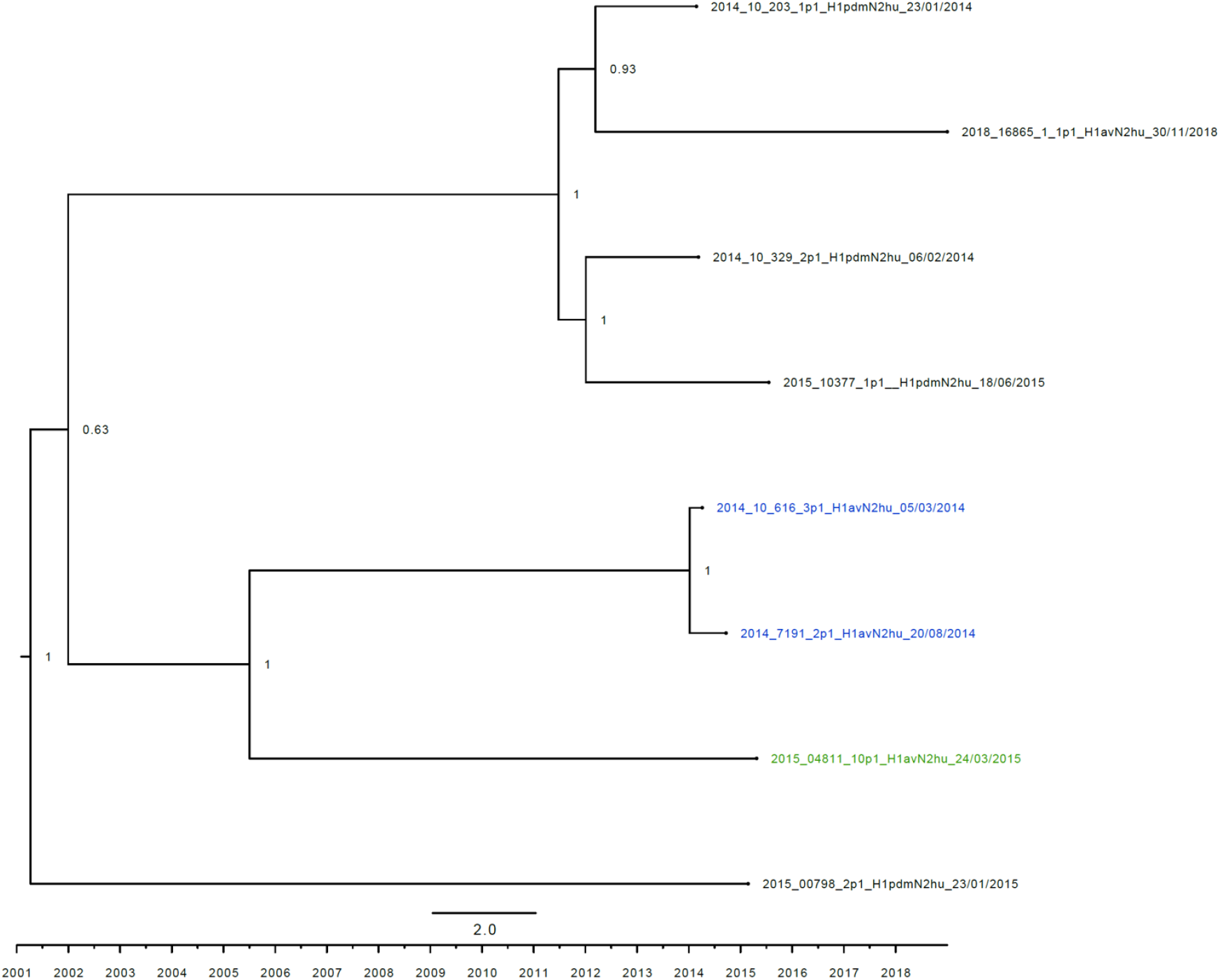
Strict molecular clock tree of the N2hu sequences The x-axis indicates the time in years and each tick indicates half a year. A blue taxon indicates that the sample carried an internal gene cassette of avian origin, whereas a green taxon indicates that the sample carried a partial internal gene cassette of H1N1pdm09 origin. A black taxon indicates that the sample carried an internal gene cassette of H1N1pdm09 origin.

**Supplementary figure 8.**
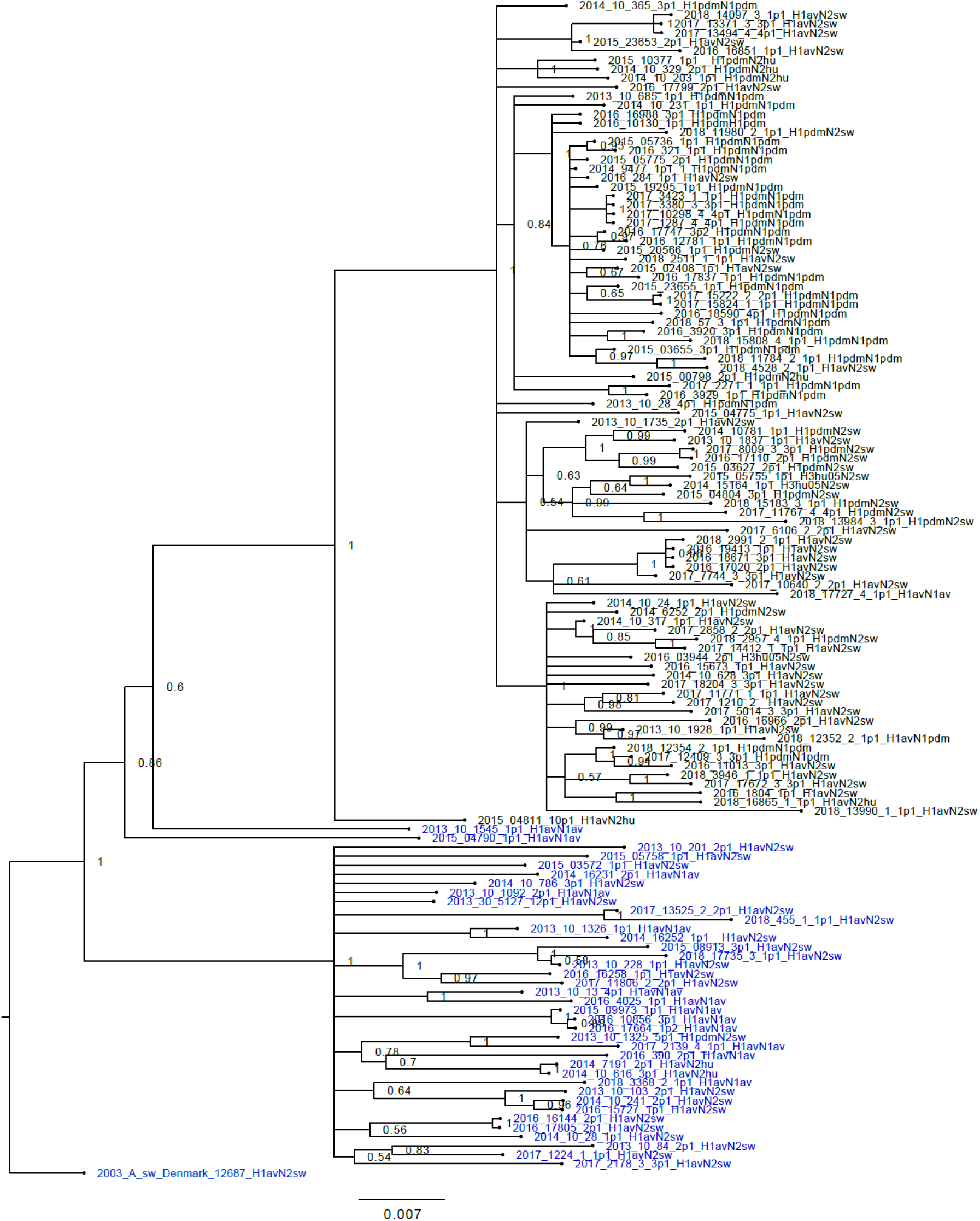
Bayesian phylogenetic tree of the M sequences “2003_A_sw_Denmark_12687” was used as the outgroup. A blue taxon indicates that the M gene of the sample is of avian-like origin, whereas the a black taxon indicates that the M gene of the sample is of H1N1pmd09 origin.

**Supplementary figure 9.**
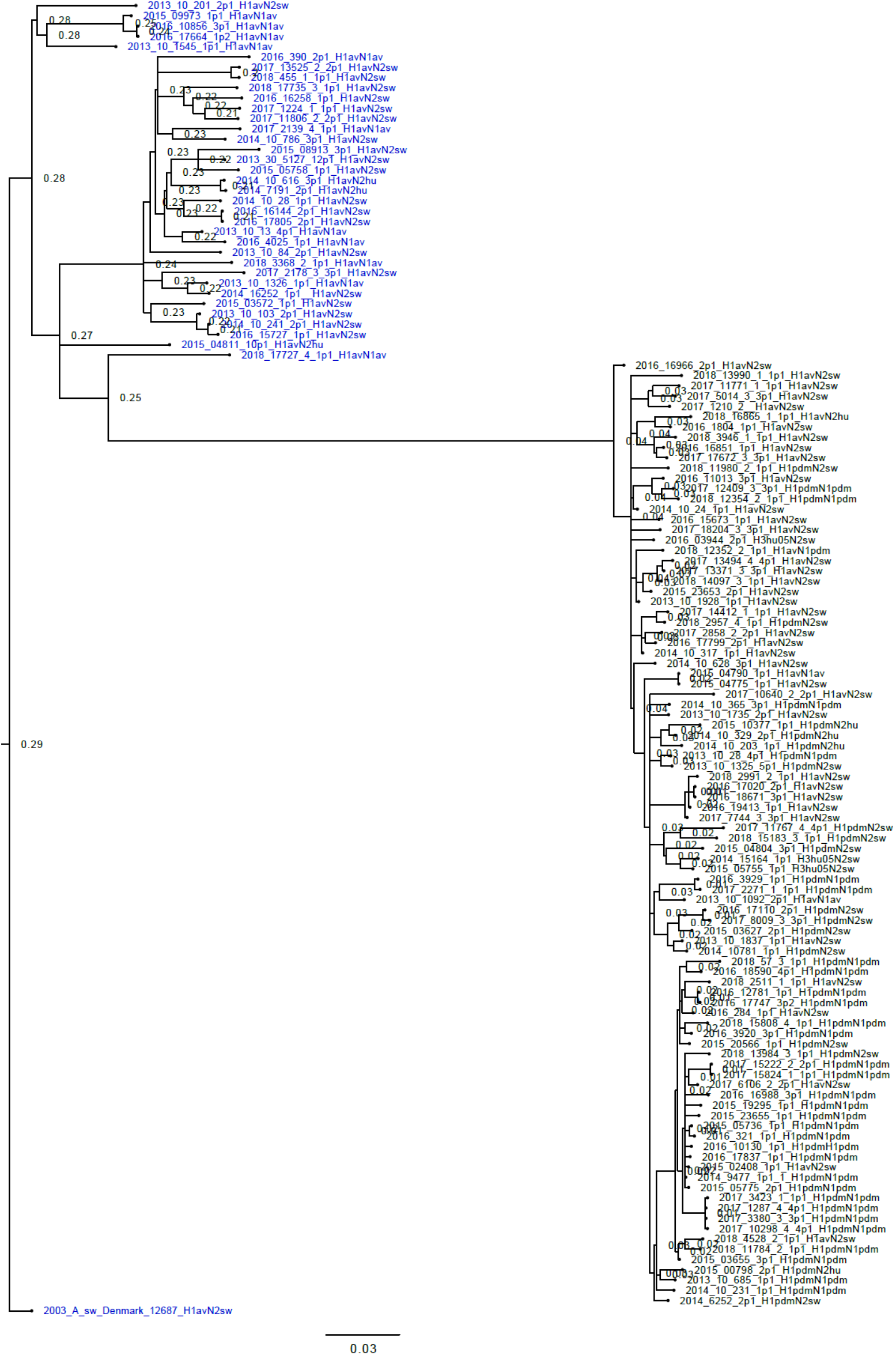
Bayesian phylogenetic tree of the NP sequences “2003_A_sw_Denmark_12687” was used as the outgroup. A blue taxon indicates that the NP gene of the sample is of avian-like origin, whereas the a black taxon indicates that the NP gene of the sample is of H1N1pmd09 origin.

**Supplementary figure 10.**
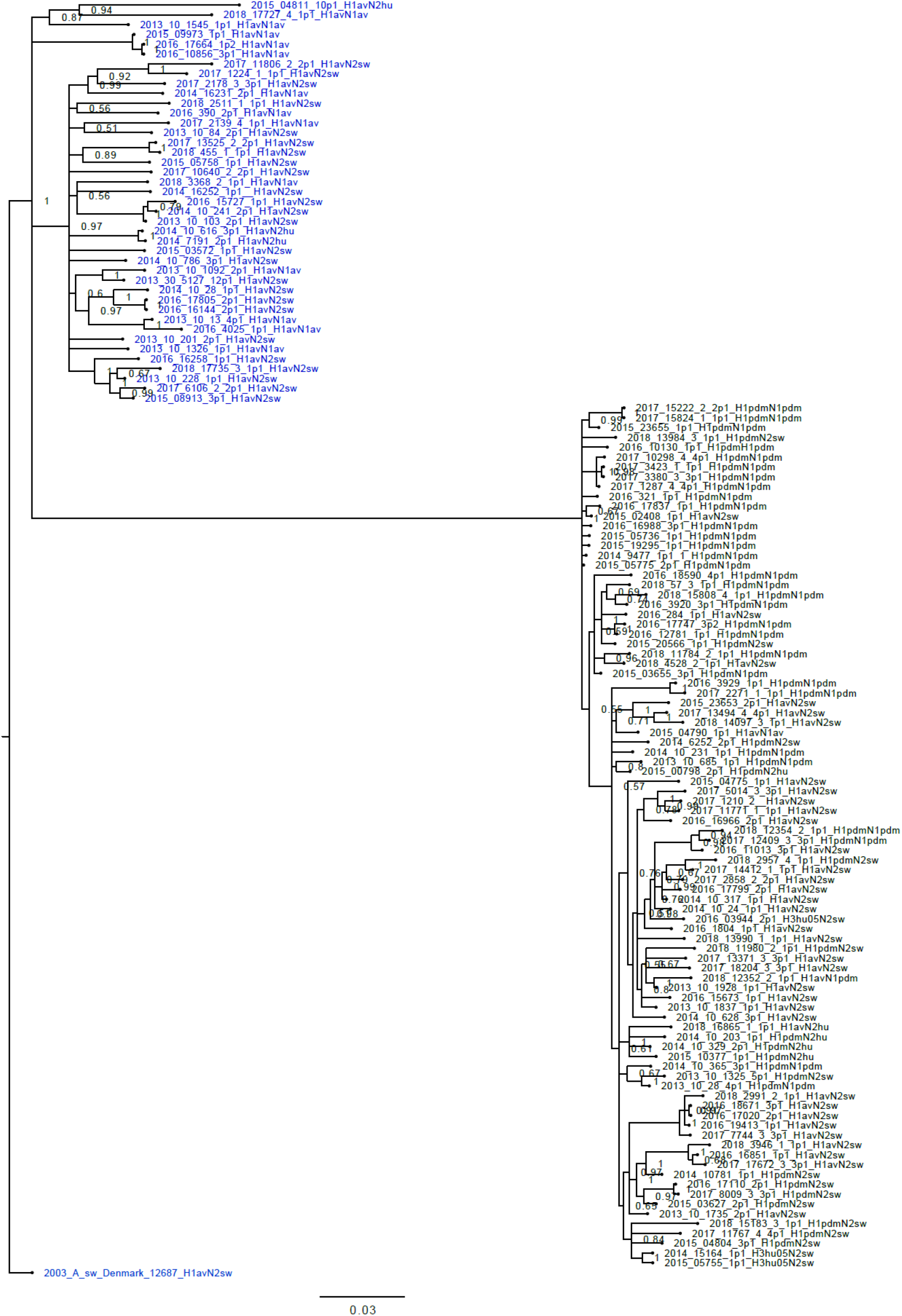
Bayesian phylogenetic tree of the NS sequences “2003_A_sw_Denmark_12687” was used as the outgroup. A blue taxon indicates that the NS gene of the sample is of avian-like origin, whereas the a black taxon indicates that the NS gene of the sample is of H1N1pmd09 origin.

**Supplementary figure 11.**
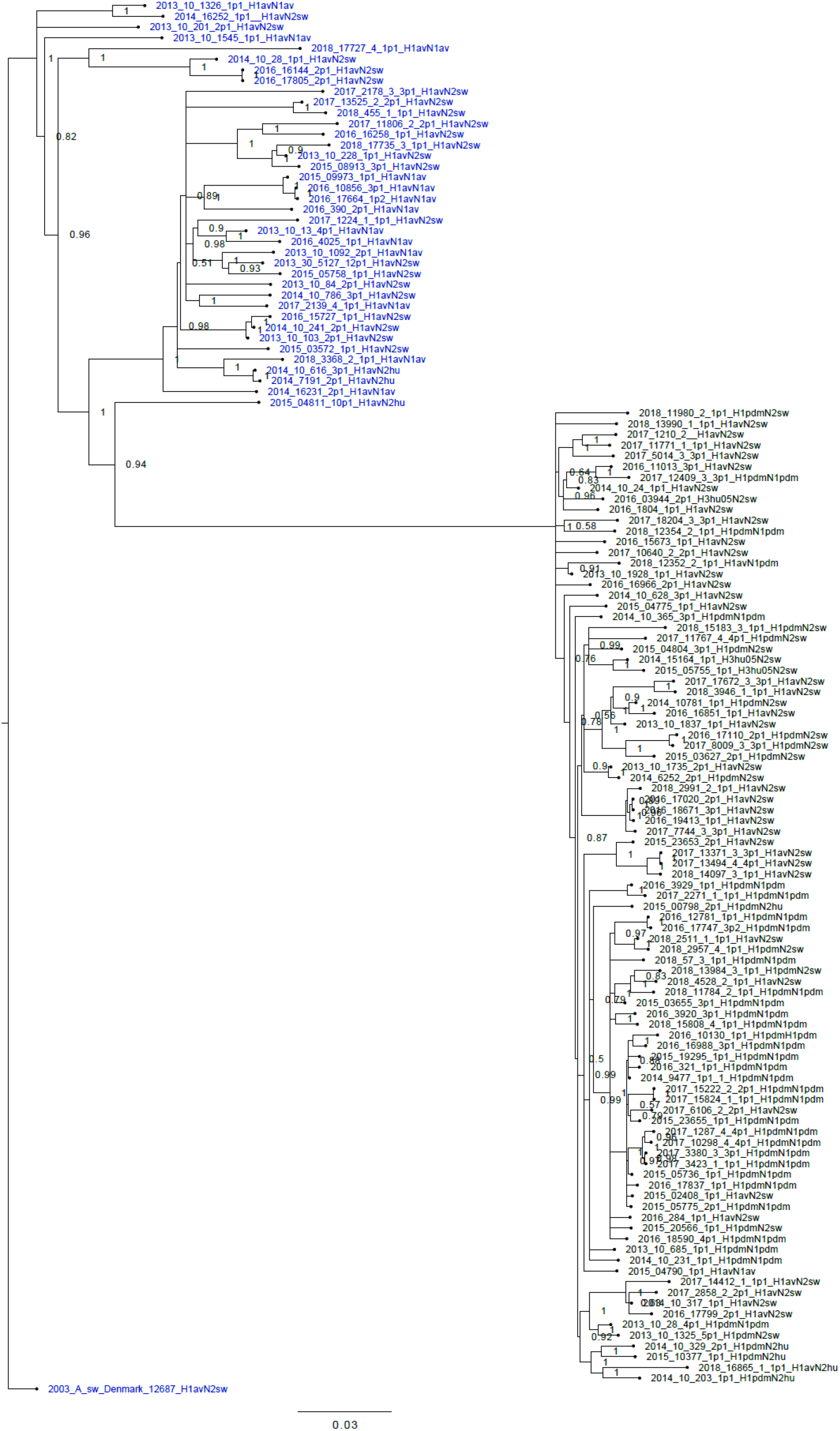
Bayesian phylogenetic tree of the PA sequences “2003_A_sw_Denmark_12687” was used as the outgroup. A blue taxon indicates that the PA gene of the sample is of avian-like origin, whereas the a black taxon indicates that the PA gene of the sample is of H1N1pmd09 origin.

**Supplementary figure 12.**
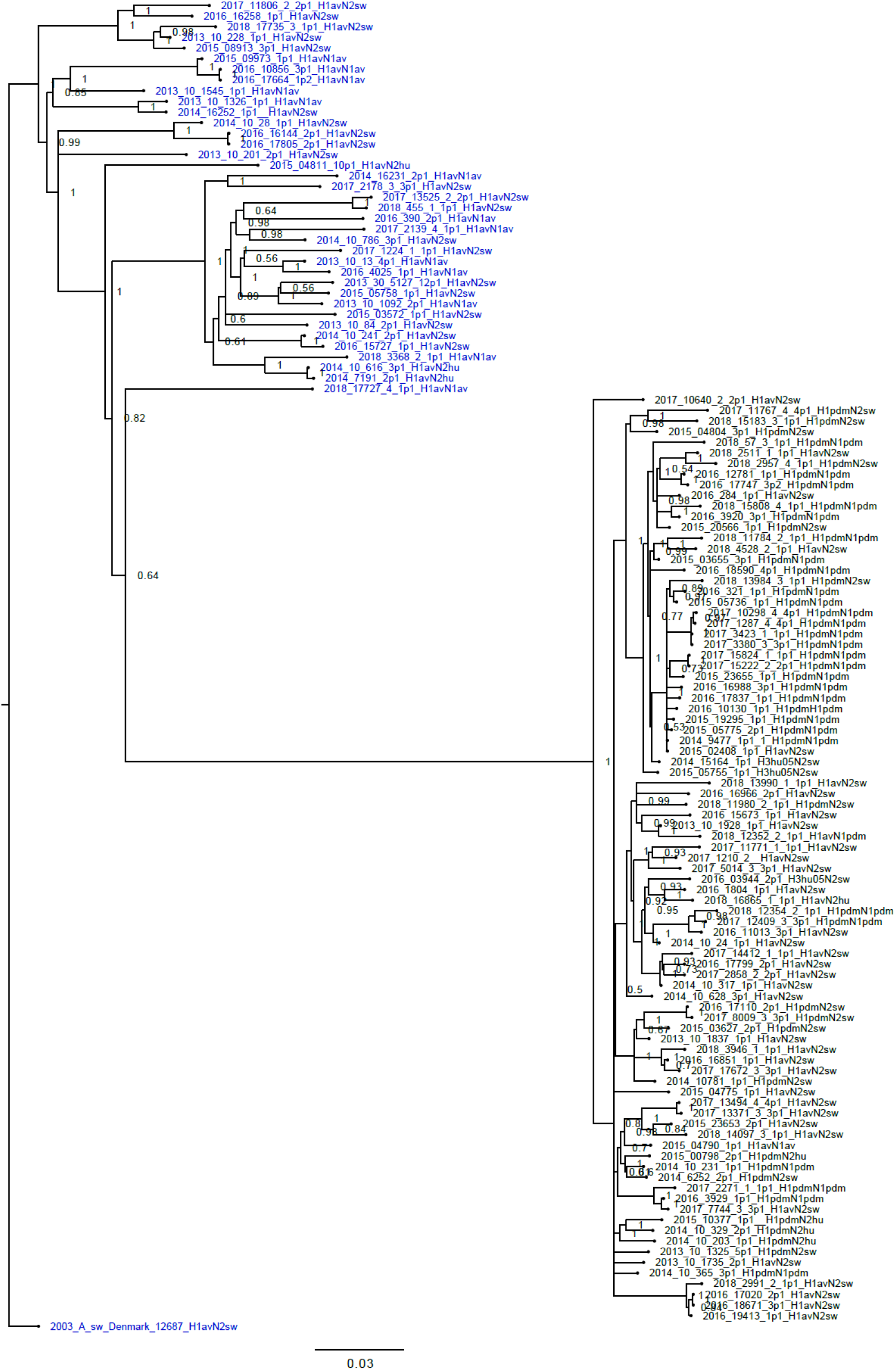
Bayesian phylogenetic tree of the PB1 sequences “2003_A_sw_Denmark_12687” was used as the outgroup. A blue taxon indicates that the PB1 gene of the sample is of avian-like origin, whereas the a black taxon indicates that the PB1 gene of the sample is of H1N1pmd09 origin.

**Supplementary figure 13.**
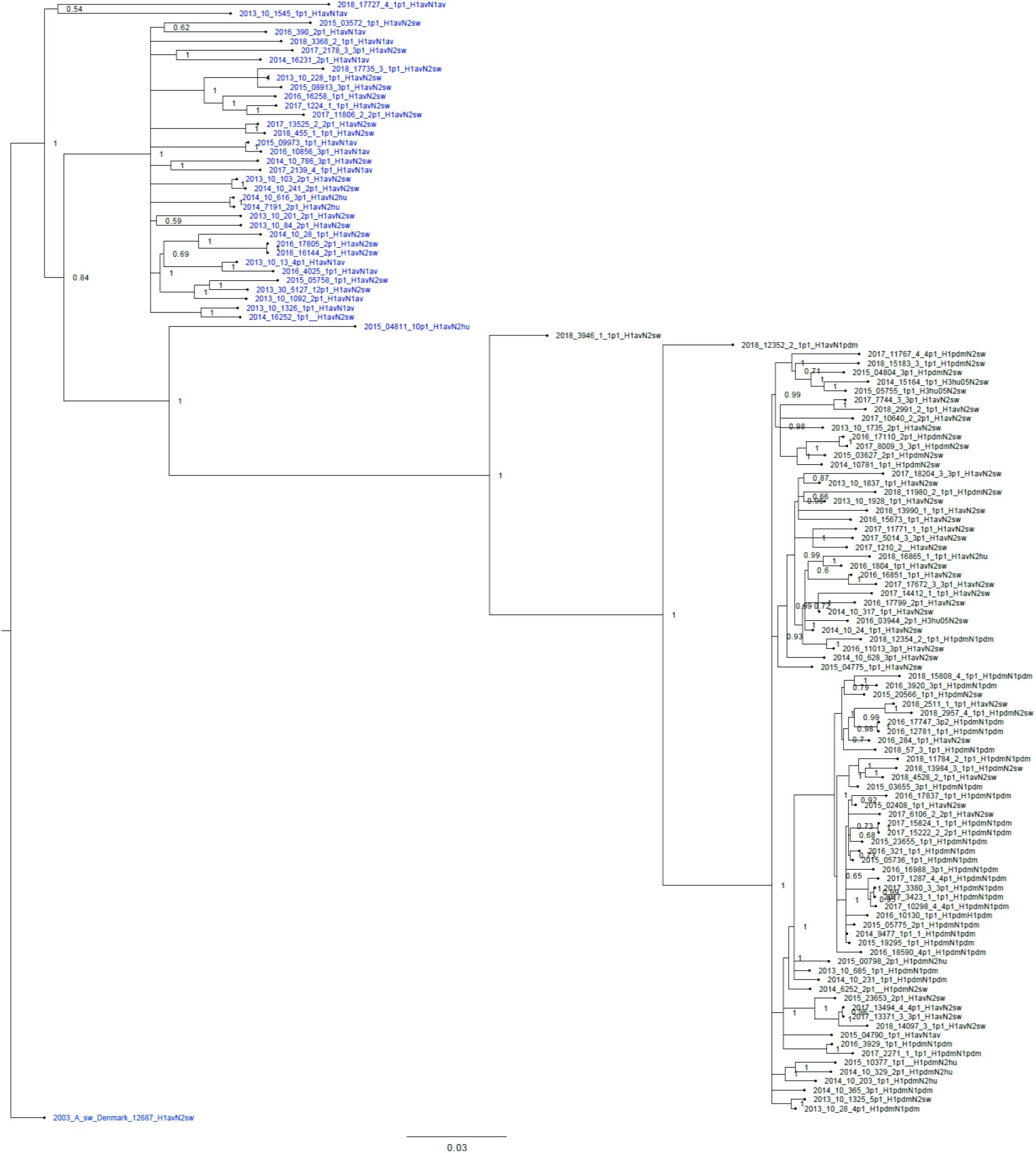
Bayesian phylogenetic tree of the PB2 sequences “2003_A_sw_Denmark_12687” was used as the outgroup. A blue taxon indicates that the PB2 gene of the sample is of avian-like origin, whereas the a black taxon indicates that the PB2 gene of the sample is of H1N1pmd09 origin.

